# A mouse model of cardiac AL amyloidosis unveils mechanisms of tissue accumulation and toxicity of amyloid fibrils

**DOI:** 10.1101/2024.07.18.604040

**Authors:** G Martinez-Rivas, MV Ayala, S Bender, GR Codo, K Swiderska, A Lampis, L Pedroza, M Merdanovic, P Sicard, E Pinault, L Richard, F Lavatelli, S Giorgetti, D. Canetti, A Rinsant, S Kaaki, C Ory, C Oblet, J Pollet, E Naser, A Carpinteiro, M Roussel, V Javaugue, A Jaccard, A Bonaud, L Delpy, M Ehrmann, F Bridoux, C Sirac

**Author notes:** Correspondence: Christophe Sirac, BioPIC team, UMR CNRS 7276 INSERM 1262, CBRS room 110, 2 rue du Pr Descottes 87000 Limoges, France;.

## Abstract

AL amyloidosis is one of the most common types of systemic amyloidosis, caused by the deposition in tissues of fibrillar aggregates of abnormal immunoglobulin (Ig) light chain (LC), leading to organ dysfunction. The most frequent and severe forms affect the kidneys and heart, the latter being associated with a poor prognosis. Despite extensive efforts to decipher the mechanisms of fibril formation and their toxicity, the lack of reliable in vivo models hinders the study of the disease in its physiological context. We developped a transgenic mouse model producing high amounts of a human AL light chain (LC). While mice exceptionnaly develop spontaneous AL amyloidosis and do not exhibit organ toxicity due to the circulating amyloidogenic free LC, a single injection of amyloid fibrils, made up of the variable domain (VL) of the human LC, or soluble VL led to amyloid deposits in the heart, vessels, spleen and, to a lesser extent, in the kidney and other visceral tissues. AL fibrils in mice contain both full length and fragmented LC with a fragmentation pattern highly superposable to that of human AL fibrils from the same LC subgroup (IGLV6-57). They also develop an early cardiac dysfunction closely resembling the human disease with increased NT-proBNP,and activation of pathways involved in the extracellular matrix remodeling and fibrosis. Overall, this transgenic AL model closely reproduces human cardiac AL amyloidosis and shares with humans the biochemical composition of the deposits, arguing for a conserved mechanism of amyloid fibrils formation. It also shows that a partial degradation of the LC is likely required to initiate amyloid fibril formations. This model offers a new avenue for research on AL amyloidosis and fills an important gap for the preclinical evaluation of new therapies.

## INTRODUCTION

AL amyloidosis, one of the most severe and frequent form of systemic amyloidosis, is characterized by the deposition of amyloid fibrils constituted by a monoclonal immunoglobulin free light chains (LCs) produced in excess by a B or plasma cell clone (*1*). The LCs involved in amyloidosis have a propensity to aggregate into the characteristic β-sheet structure of amyloid fibrils, and accumulate in the extracellular compartments of tissues, leading to organ dysfunction. Renal and cardiac manifestations are the most frequent, the latter being associated with poor outcomes(*2, 3*).

The molecular mechanisms that lead to the aggregation of LCs have been extensively studied, primarily *in vitro* due to the lack of other models reproducing the early stages of the disease. The variable domain (VL) of LCs has long been suspected to be the causative part of AL amyloid fibrils formation since only few VL germline genes account for most of the cases (*4–7*) and destabilizing mutations acquired during affinity maturation in the V domain has been shown critical for amyloidogenicity *in vitro* (*8–10*). Consequently, our knowledge on LC aggregation process has been mostly obtained from isolated amyloidogenic VL since full-length LCs seem to be resistant to aggregation under physiological conditions (*11, 10, 12*). Accordingly, the recent 3D resolution of *ex vivo* AL amyloid fibrils with cryo-electron microscopy (CryoEM) confirmed that the cross β-structured interactions within the core of the fibrils are primarily established by the VL domains (*13, 14*). The remaining portion of the LC, the constant domain (CL), seems to be disorganized outside the fibrillar structure and partially cleaved. Several proteolytic points have been identified primarily in the N-terminal part of the VL and throughout the CL (*15, 16*). These cleavage sites correspond to the outer parts of the LCs, not protected by the fibril-core interactions. This prompts the question of whether proteolysis in these LCs occurs before or after their aggregation. The multiple cleavage pattern in sites not accessible in the native dimers suggest a fragmentation subsequent to aggregation (*16*). But the high amyloidogenicity of some fragmented species suggests that proteolysis of the LCs could also be required to initiate amyloidosis formation (*12*). This question highlights a fundamental gap in our understanding of the pathogenesis of AL amyloidosis.

In addition to the assumed effects of amyloid fibril accumulation in tissues, the soluble form of LCs may also contribute to cardiac toxicity. Patients responding to treatments which aim at reducing circulating LCs show a significant decrease in NT-proBNP concentrations, correlated with improved cardiac function, in spite of the absence of a significant decrease in the amyloid burden (*17–20*). Studies conducted *in vitro* using cardiomyocyte and cardiac fibroblast cultures, as well as in *C. elegans* and Zebrafish models, supported this theory (*21–25*). Exposure of these models to soluble amyloidogenic LCs leads to cellular stress through internalization of the LC and increased production of reactive oxygen species (ROS) and the activation of a non-canonical MAPK pathway. As a result, lysosomal dysfunction, autophagy impairment and mitochondrial damage were observed, followed by cell death. Although these studies provide valuable insight into LC toxicity on cardiac cells, none of them fully reproduces the tissue complexity of human physiology and the underlying mechanisms that occur in patients, neither the deposition and accumulation of amyloid material composed of the LCs. Translating these findings to human physiology remains challenging, and the applications of these models beyond toxicity studies are limited.

Several approaches to create rodent models of AL amyloidosis have been attempted over the years. Firsts attempts consisted in the massive injection of Bence-Jones proteins, purified fibrils from patients or the so-called “amyloidoma” consisting in a crude grinded human tissue containing amyloid material (*26–29*). Although localized amyloid material is present in both models, their application is limited to therapeutic studies since they poorly reproduce the human pathophysiology and organ involvement. Other models have used transgenic approaches to produce amyloidogenic LCs endogenously in the mice or rat with mostly disappointing results (*30–32*). One of them succeeded at reproducing amyloid deposition, even though deposits were localized in the stomach of aged mice, suggesting a destabilization of LCs in the local acidic environment which promoted aggregation rather than a physiological amyloid formation in classically involved organs (*33*). This apparent resistance to amyloidosis in mice was also observed for other types of systemic and localized amyloidosis and was attributed to a better proteostasis and a rapid turnover of proteins (*34–36*).

One of the main limits of these models is the levels of circulating LCs which are much lower than those observed in patients, likely failing to reach the threshold needed to initiate fibril formation (*37*). To overcome this limit, we have developed a unique transgenic approach which allows to achieve high levels of circulating pathological LCs into the mice. This strategy has been successful for modelling non-amyloid monoclonal LC-related deposition diseases affecting the kidneys, including light chain deposition disease (LCDD) and light chain proximal tubulopathy (LCPT) with Fanconi syndrome, allowing to reproduce the typical renal LC deposits observed in patients (*38, 39*).

In the present study, we applied this transgenic approach to create a mouse model producing an AL amyloidosis LC to overcome the lack of *in vivo* models for studying this devastating disease and provide a reliable tool for preclinical investigations of new therapies or diagnostics. Mice present with a high level of free amyloidogenic LCs but only exceptionally develop spontaneous systemic AL amyloidosis. However, upon induction not only with preformed fibrils but also with soluble amyloidogenic VL, they develop robust amyloid deposits in heart, spleen, vessels and to a lesser extent kidney and liver. This study characterizes this new AL mouse model and offers new insights into the pathophysiological mechanisms of AL amyloidosis *in vivo*.

## RESULTS

### High levels of circulating amyloidogenic lambda free light chains in mouse is not enough *per s*e to induce AL amyloidosis or symptomatic toxicity

To generate the mouse model of AL amyloidosis, we used the cDNA coding a monoclonal λ LC derived from a patient with AL amyloidosis (λS-PT). Patient FLC levels at diagnosis were 128 mg/L for λ and 18 mg/L for κ LC, associated with an IgAλ monoclonal component in serum, and 6.5% of medullar plasma cells (PC) infiltration. Clinical manifestations at diagnosis included chronic kidney disease (serum creatinine level 191 µmol/l, hypoprotidemia 56 g/l, heavy proteinuria 5.8 g/24h) and typical features of amyloid cardiomyopathy (diffuse microvoltage on ECG, interventricular septum thickness of 14 mm, diastolic dysfunction, preserved left ventricular ejection fraction at 60%). Serum NT-proBNP was initially at 13173 pg/mL, likely exacerbated by the renal dysfunction. Kidney biopsy confirmed the presence of AL amyloid deposits composed of a λ LC (**Fig. 1A**). Monoclonal LC (λS-LC) cDNA sequencing revealed an IGLV6-57-derived variable germline gene rearranged to an IGLJ3 junction segment and an IGLC3 constant domain. This V germline gene is the most frequent in AL amyloidosis and is classically associated with dominant renal and cardiac involvements (*40*). Mutation analysis revealed 11 mutations compared to the germline IGLV6-57 and IGLJ3 sequences (**Fig. S1A**). By analyzing the sequence on Aggrescan4D (*41*), three of these mutations were shown to create a hydrophobic side, resulting in an aggregation-prone region exposed to the solvent in the dimeric conformation (**Fig. S1B**).

**Figure 1:**
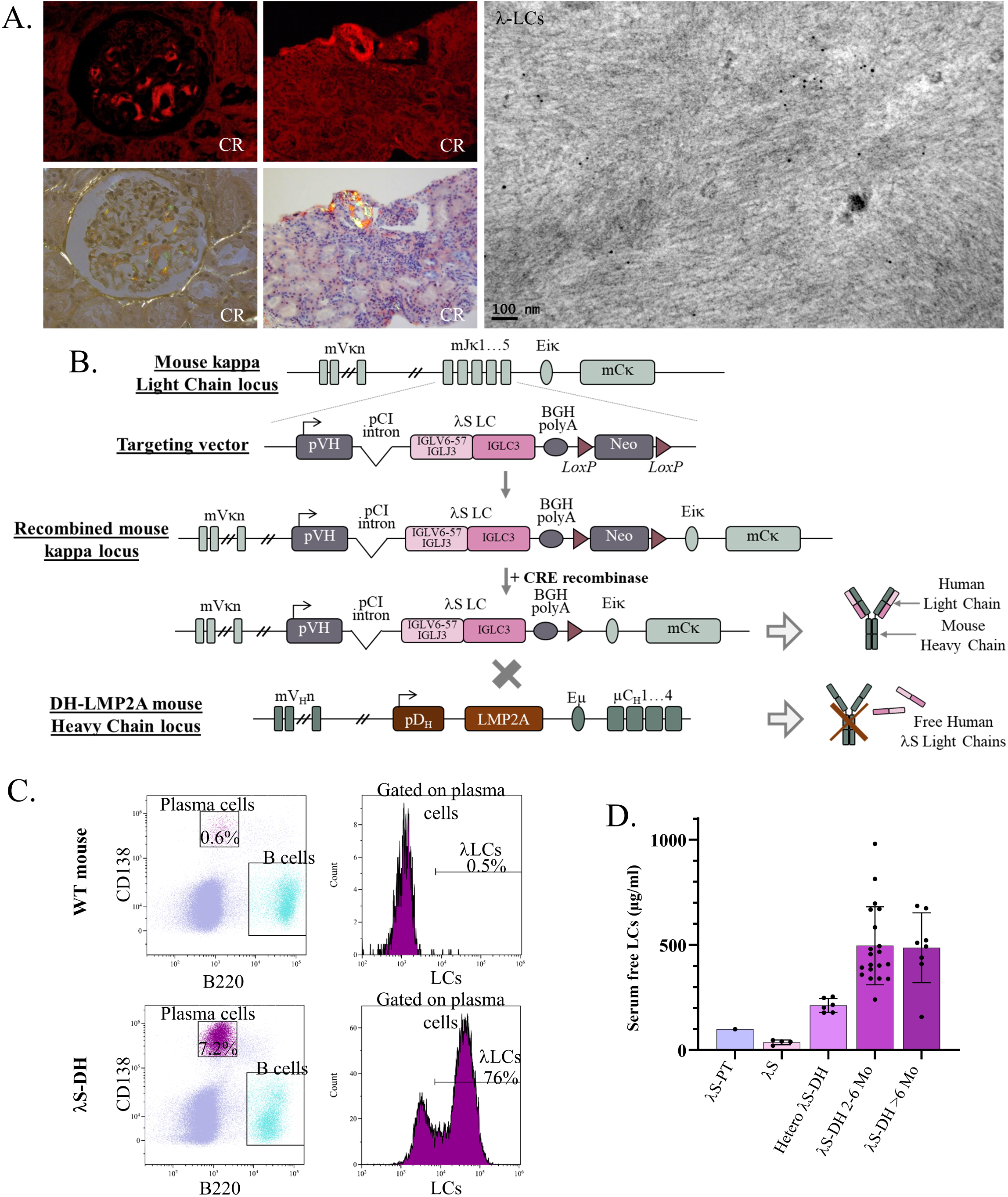
Generation of the transgenic mouse model λS-DH. A. Kidney biopsy from the patient λS (λS-PT) showing the Congo Red (CR) staining in fluorescence (top) and its birefringence under polarized light (bottom) in a glomerulus and in a blood vessel. The characterization of the deposits was performed by immunoelectron microscopy with anti λ-LCs gold-labeled antibody (right). B. The transgenic strategy used to achieve a high production of human free light chains in the mice consists in the insertion of the λS-LC from the λS-PT in the mouse κLC locus. These mice were backcrossed with the DH-LMP2A mice to avoid the association between human LCs and mouse heavy chains C. Analysis of plasma cells in the spleen of λS-DH mice by flow cytometry. CD138 and B220 staining allows to differentiate plasma cells from B cells and all other splenic cells. Staining of hλ-LC allows to show the plasma cells producing the transgenic λS-LC. Representative examples from the analyzed mice (n=3 WT and n=3 λS-DH) D. Serum dosage of the circulating free light chains in the patient at diagnosis (λS-PT) and the transgenic mice without (λS) and with (λS-DH) the DH-LMP2A allele. The histograms correspond to the mean of the group and the error bars to the standard deviation (SD).

High production of the pathogenic human free λS-LC in the mouse was achieved thanks to a modified method derived from our previously published immunoglobulin (Ig) kappa knock-in strategy (see methods) (*39*). Briefly, we targeted the mouse kappa locus to introduce the complete cDNA coding the human amyloidogenic λS-LC sequence, replacing the IGKJ segments. Then we crossed these mice with the DH-LMP2A strain (thereafter called DH) (*42*) that allows a normal B cell development with increased plasma cell number in the absence of Ig heavy chains (**Fig. 1B**) (*43*). These mice produce in the serum approximatively 100 µg/mL of polyclonal mouse κ LCs and there were used as controls thorough the study except when otherwise stated. As expected (*43, 39*), the resulting double homozygous λS-DH mice presented with a high number of spleen PCs (8.9% +/- 3.9, n = 3) mostly producing the human transgenic λS LC (76.3% +/- 2.7), mimicking features of monoclonal gammopathies (**Fig. 1C**).

FLC serum level reached 493.0 ± 33.4 µg/ml (mean ± SEM, n = 28) in 6-8 month-old λS-DH mice, which is 4 fold more than in the patient λS-PT at diagnosis (**Fig. 1D**). This FLC level is all the more elevated as the serum half-life of the human λS-LC is extremely short in mice (~10 minutes) as determined by recombinant λS-LC injections in WT C57/B6 mice (**Fig. S1C**). Serum FLC level in the λS mice (without DH backcrossing) was only 37.2 µg/ml ± 5.6 (mean ± SEM, n=4), likely due to the association of the human LC to mouse HC, forming full Igs (*38*). Western blot using a mouse anti-human IGLV6 antibody confirmed the presence of the human λS-LC in the sera of λS-DH mice (**Fig**. **S1D**).

As expected, human LCs were readily detectable by immunofluorescence in the spleen, especially in extrafollicular plasma cells (bright green staining) and in proximal tubular cells of the kidney, corresponding to the physiological tubular reabsorption of the LC (**Fig. S1D**) However, we did not observe any amyloid deposits in 6-8 month-old mice (n=10). λS-DH mice had a normal lifespan compared to control DH mice with no signs of morbidity. Serum creatinine and albuminuria levels at 8 months old were 4.40 μM/L ± 1.51 (mean ± SEM, n=5), and 43.93 µg/ml ± 9.54 (n=6), respectively, which is comparable to previously published data in DH and WT mice (*39*). Altogether, these results show that the transgenic amyloidogenic LC do not seem to display apparent direct toxicity or have a global negative effect on the survival of mice.

### Variable domain but not full length of the **λ**S-LC is amyloidogenic *in vitro*

In order to understand why λS-DH mice do not develop AL amyloidosis, we sought to characterize the amyloidogenicity of the λS-LC *in vitro*. Since most studies on *in vitro* amyloidogenicity of LCs were conducted on VL fragments alone and full length LCs were shown to poorly aggregate under physiological conditions (*11, 12*), we produced both recombinant forms, namely the Full-Length LC (rλS-LC) and its variable domain alone (rλS-VL). As expected, rλS-LC mainly constituted dimers that were stabilized by the disulfide bond between constant regions and rλS-VL remained under its monomeric form (**Fig. S2A and Fig. S2B**).

We studied the stability of rλS-VL and rλS-LC by their thermal denaturation over a temperature range from 35 to 95°C (**Fig. 2A**). The T_m_ of the rλS-VL protein was 38.7°C, which is similar to the temperature of Wil, another amyloidogenic IGLV6-57–derived VL widely used for *in vitro* aggregation studies (*44*) althoughthey shared only 80.0% sequence similarities (**Fig. S1A**). Conversely, the rλS-LC protein had two T_m_ at 48.5 and 59.4, likely corresponding to the minor monomeric form and the dimeric form respectively. The thermal unfolding midpoints correlated with the ability of these two rλS proteins to start fibrillogenesis *in vitro*, as observed in Thioflavin T assays (ThT) (**Fig. 2B**). The rλS-VL rapidly resulted in the increase of ThT fluorescence, starting at 2 days under aggregating conditions, while the rλS-LC never increased throughout the entire experiment (96 h). Electron microscopy confirmed the presence of fibrils formed by the rλS-VL while no fibrillar structures were detected with the rλS-LC (**Fig. 2C**).

**Figure 2:**
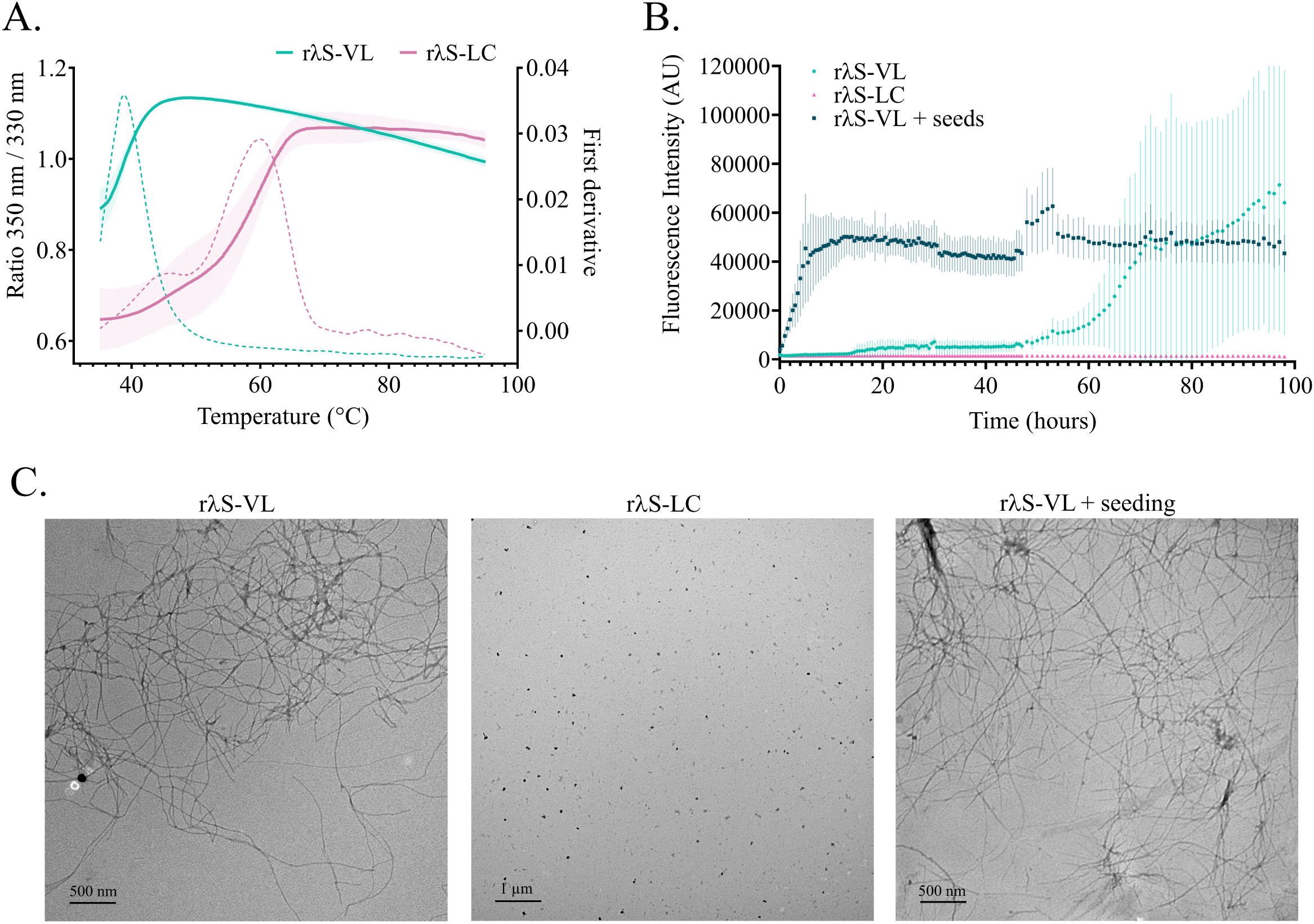
*In vitro* characteristics of λS-LC species. A. Stability curves of the recombinant λS proteins, rλS-LC and rλS-VL, was measured by the ratio of the fluorescence measured at 350 and 330 nm (solid line) upon thermal denaturation. The melting temperature (Tm) for each sample was given by the maximum peaks of the 350/330 nm ratio first derivative (dotted line). The curves correspond to the mean of 3 different measurements for the rλS-LC and 4 for the rλS-VL, and the error bars correspond to the SD. B. Kinetics of fibril formation *in vitro* with the different λS protein species at 37°C and 300rpm in aggregation buffer and with (rλS-VL+seeds) or without seeds (rλS-VL and rλS-LC) and in presence of 2 µM of Thioflavin T (ThT). The aggregation kinetic was followed by the measurement of the fluorescence of ThT (ex: 448 ± 10 nm; em: 482 ± 10 nm). The curves correspond to the mean of at least 3 different experiments in duplicate. The error bars correspond to the SD. C. TEM analysis of the samples from (C) after 60h for the rλS-VL (left) and rλS-LC (middle), and 16h for the rλS-VL+seeds (right), to confirm the presence of fibrils in the samples. Images are representative examples of each one of the samples.

We further tried to accelerate the kinetics of fibrillogenesis *in vitro* using a seeding with sonicated preformed fibrils. Preformed fibrils can act as a platform to bypass the nucleation phase of amyloidogenic proteins (*37*). In this aim, we sonicated the fibrils obtained with the protein rλS-VL and mix them together with soluble rλS-VL. This highly increased the kinetic of fibrillogenesis of rλS-VL, reaching the plateau phase after a few hours (**Fig. 2B and Fig. 2C**). Altogether, these results confirm that the full-length λS-LC seems to be resistant to fibril formation *in vitro*, which corroborates the *in vivo* findings in the λS-DH mice. However, we found a high amyloidogenicity of the λS variable domain.

### Seeding with recombinant **λ**S-VL fibrils induces fast AL amyloid deposition *in vivo*

Seeding with Amyloid Enhancing Factor (AEF), basically a crude grinded tissue containing amyloid, or with purified *ex vivo* fibrils from a human or mouse source was necessary to induce amyloidosis formation in other animal models of systemic amyloidosis, including AA and ATTR models (*45–47*). Unfortunately, in our case, we do not have access to tissue from the patient (λS-PT) but we hypothesized that the rλS-VL fibrils made *in vitro* could serve as seeds for the nucleation and elongation of amyloid fibrils with the endogenous λS-LC in our transgenic mice (**Fig. 3A**). Accordingly, the intravenous injection of *in vitro* rλS-VL sonicated fibrils resulted in the formation of amyloid deposits in a systemic manner, starting as soon as 48h after the injection. Congo red birefringence and fluorescence showed that cardiac and surrounding vascular tissues were principally affected together with the spleen (**Fig. 3B**). In contrast with the patient from whom the LC was extracted (λS-PT), λS-DH mice rarely developed kidney deposits, only visible in mice with high deposits in heart and spleen (**Fig. 3B**). Nevertheless, deposits in the kidney were localized in the glomeruli, as commonly observed in AL patients and in the patient λS-PT. Cardiac deposits were found mainly in the myocardial compartment of the ventricular and atrial walls, along the muscle fibers, resulting in diffuse thin filaments outlining the cells (**Fig. 3C**). In cardiac blood vessels, deposits were localized throughout the tunica media, but also in the adventitia (**Fig. 3C**). Not all seeded mice developed amyloidosis but the penetrance increase with time, ranging from 28 % one week after injection to 100 % after 6 months (**Fig. 3D**). Since the variability on the amyloid burden and time of appearance was significant, a score system was established according to the Congo Red staining in the heart (**Fig. S3A**). Using this score, we showed that there is a high heterogeneity of amyloid burden in the positive mice whatever the time after induction is, with the lower average score in short term induction mice (< 1 week, mean score 1,50, range=1-3, n=10) and higher score in long term amyloidosis induction (> 6 months; mean score 2,67, range=1-3, n=6). However, except for long-term induction, we did not observe significant increase in amyloid burden with time (**Fig. 3E**). We did not detect amyloid deposits in control mice injected with the same amount of *in vitro* fibrils, and either producing a human monoclonal non-amyloidogenic κLC (κR-DH, n=11), polyclonal murine LCs (DH, n=7), or full polyclonal Ig (WT, n=9) (**Fig. 3D and Fig. S3B**). This result ruled out the possibility that amyloid deposits detected in organs were due to the injected fibrils themselves. Ultrastructural studies of the cardiac deposits showed the typical conformation of amyloid fibrils, as unbranched, long filaments intertwined in the extracellular compartment of the tissues (**Fig. 3F**). In high score mice, we observed myocardial interstitial fibrosis, a typical observation in patients **(Fig. 3G)** (*48*). We further confirmed the typing of amyloidosis in our mice by immunofluorescence with an anti-human λ-LC antibody. For that purpose, we selected a monoclonal antibody recognizing specifically the constant region of the human λ-LCs to ensure that amyloid fibrils are made up of the transgenic full length λS-LC and not only the injected VL fibrils. Accordingly, we detected a strong staining for λ-LCs in the positive mice, which mostly colocalized with CR positive areas in heart **(Fig. 4A),** spleen and kidney **(Fig. 4B)**. Interestingly, in hearts with low amyloid burden (score Low), we readily detected human λ-LCs staining that not only colocalized with CR clusters but also stained CR negative intercellular spaces (**Fig. S3C**). Such staining was never detectable in non-induced mice or control induced mice (**Fig. S3B**). This suggests that prefibrillar aggregates or non-mature amyloid fibrils, not yet detectable by CR staining, are likely present in the tissue. Systemic AL deposits were also found in different amounts in the tongue, liver, lung and fat (**Fig. S4**). Immuno-electron microscopy confirmed the typing, showing strong binding of the gold-labeled anti-human λ-LC to amyloid fibrils but not with anti-human κ-LC in the cardiac tissue of λS-DH mice (n=2) **(Fig. 4C)**. Altogether, these results show that the transgenic full length λS-LCs are the main constituent of amyloid fibrils in the λS-DH mice and can therefore elongate amyloid seeds composed of the VL alone to form mature amyloid deposits.

**Figure 3:**
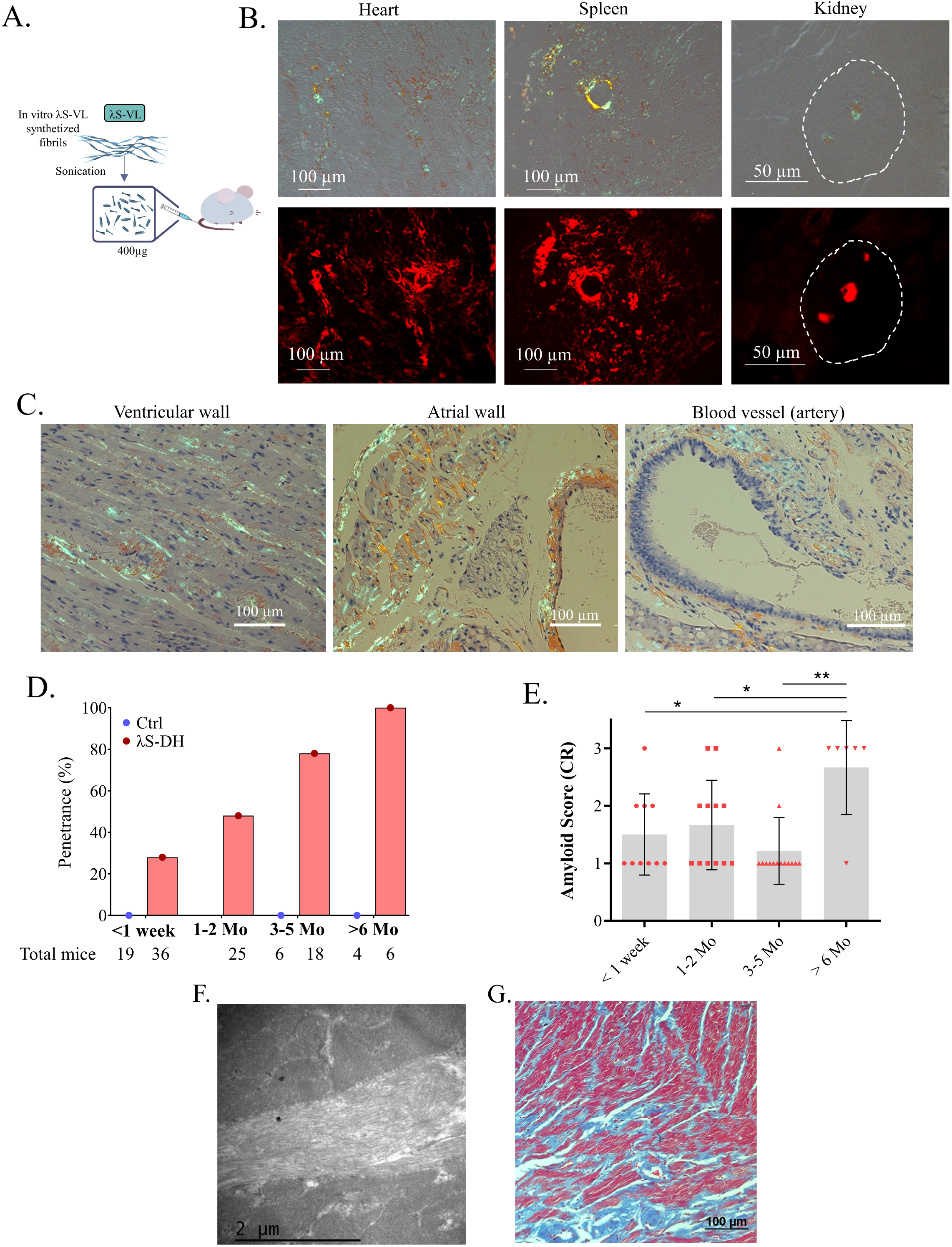
Amyloid deposits induction in λS-DH mice with VL fibrils. A. Protocol of amyloidosis induction with VL fibrils. B. Histological characterization on frozen tissues of mice deposits with CR staining and analysis of the birefringence characteristic of amyloid deposits under polarized light (top) and their fluorescence (bottom) in the heart, the spleen and in the kidneys, localized in the glomeruli (white circle) of mice. Example of a mouse analyzed at 6 months after induction (high score). C. Hematoxylin-Eosin staining coupled to the Congo Red staining in a paraffin-embedded heart. Birefringence of the Congo Red was visible in the myocardium of the ventricular wall (left), but also the atrial wall (middle) and around blood vessels (right). This mouse was analyzed at 9 months after induction (high score) D. The organs from induced mice (λS-DH and controls) were analyzed histologically with a Congo Red staining at different timepoints after the injection. The penetrance of the deposits was calculated by the presence of Congo Red staining in the heart of each timepoint. The number of mice analyzed at each timepoint are indicated under the graph. E. The amyloidosis score was calculated for the positive mice by the Congo Red fluorescence in the cardiac tissue, corresponding to: Low (score 1), Mid (score 2) and High (score 3). Scores are described in the supplementary data and material and methods. The bar graphs correspond to the mean of score, and the error bars to the standard deviation. F. Electron microscopy of the cardiac tissue showing the amyloid fibrils in the extracellular compartment in a high score mouse 7 months after induction. G. Fibrotic tissue revealed with a Masson’s Trichrome staining in the heart of λS-DH mice. Example from a high score λS-DH mouse 9 months after induction.

**Figure 4:**
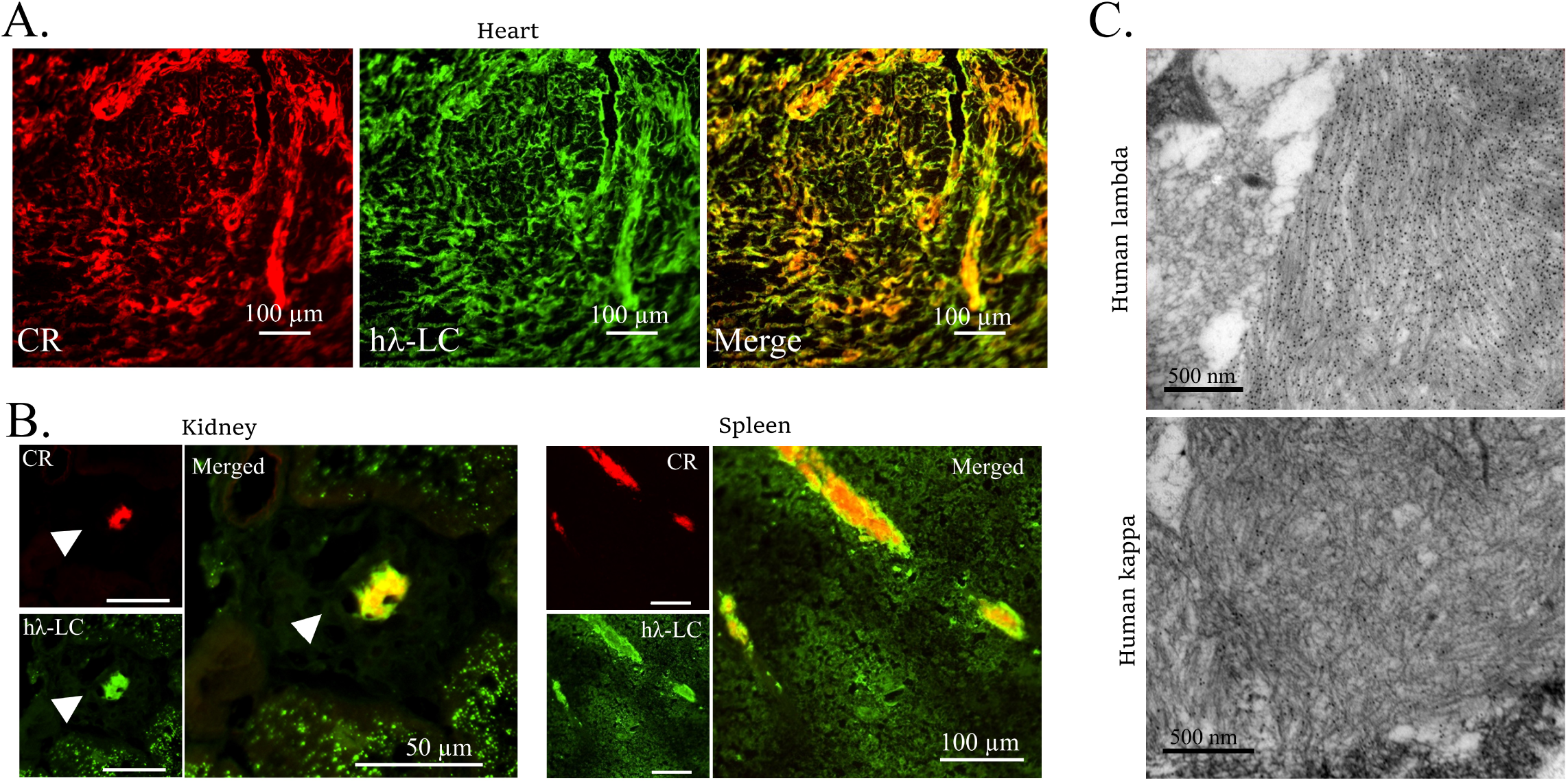
Typing of the amyloid deposits in λS-DH mice. A. Cardiac fluorescence staining of deposits with Congo Red and an anti-human λLCs antibody coupled to FITC. The merge shows the colocalization between the CR and the anti hλ-LCs staining. The anti hλ-LCs antibody used for this staining was a monoclonal antibody recognizing an epitope of the constant domain of λ-LCs. Example of a high score mouse 4 months after induction. B. Colocalization between the CR staining and the human λ-LCs was also confirmed in the kidney and the spleen of mice. Same mouse as in A. C. Confirmation of the typing of amyloidosis by immunoelectron microscopy with anti-human λ-LCs gold-labeled antibody (top) in the cardiac tissue of a high score mouse 7 months after induction. Anti-human κ LCs gold-labelled antibody was used as a control (bottom).

### Soluble amyloidogenic fragment of the **λ**S-LC is able to induce amyloid deposition in **λ**S-DH mice

Having shown that mice can develop AL amyloidosis upon seeding with AL fibrils provided the sufficient production of an amyloidogenic full length LC, we sought to determine if the native soluble variable domains of the λS light chain, shown to be the amyloidogenic part of the LC in vitro, would also be able to initiate new fibrils *in vivo.* We injected the rλS-VL fragment previously used for *in vitro* fibrils formation. To rule out the possibility that rλS-VL samples already contained aggregates prior to injection, we stored them in pure sterile Hepes buffer without NaCl and conducted several experiments including ThT assays, gel electrophoresis and HPLC that all argued for the absence of detectable aggregates (**Fig. S2B**). We injected intravenously 800 µg of soluble rλS-VL to the mice in a single dose (**Fig. 5A**). In the absence of dosage methods specific for Ig VLs, we did not evaluate the serum half-life of the soluble rλS-VL fragments. However, due to its small size (~12 kDa) and data about full length λS-LC half-life (**Fig. S1C**), we can rationally argue that the serum half-life of the soluble rλS-VL was very short. However, using this method, mice presented with amyloid deposits starting at 2 months after the injection in few mice with increased penetrance at later time points (**Fig. 5B**). Similarly to fibrils injections, the penetrance did not reach 100% and there was a heterogeneity in amyloid scores even 6 months after the initial injection of VLs. However, the general pattern of deposits and the organ tropism was the same as with fibrils injection (**Fig. 5C**). Although these results indicate that the rλS-VL fragments are likely initiating the first fibrillar aggregates, typing with the anti-λ constant region antibody confirmed that the soluble full-length transgenic LCs participated in their elongation and maturation (**Fig. 5D**). As expected, none of the control mice developed amyloidosis at 4 (n=4) or 6 (n=4) months post-injection (**Fig. 5B**). Altogether, these data demonstrate that few unstable fragments of the LCs can rapidly form stable aggregates *in vivo* that remain in the tissues and are sufficient to initiate elongation of AL amyloid fibrils.

**Figure 5:**
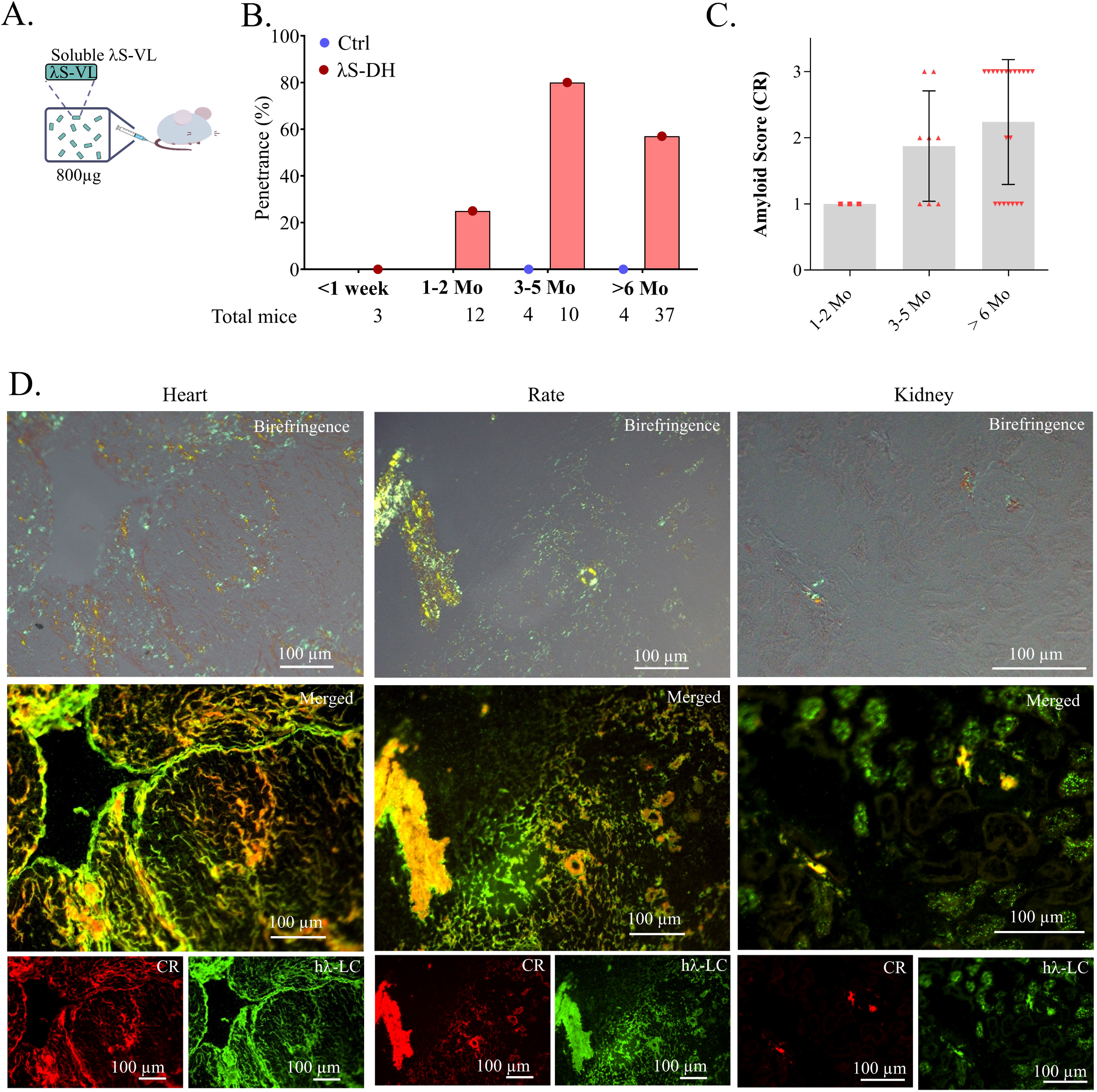
Amyloid deposits induction in λS-DH mice with soluble λS-VL VL. A. Protocol of amyloidosis induction with soluble rλS-VL B. The organs from induced mice (λS-DH and controls) were analyzed histologically with a CR staining at different timepoints after the injection, and the penetrance was calculated for each timepoint according to the presence of CR staining in the heart. The number of mice analyzed at each timepoint are indicated under the graph. C. The amyloidosis score (low = 1, mid = 2 and high = 3) was calculated for the positive mice by the Congo Red fluorescence in the cardiac tissue. The bar graphs correspond to the mean of score, and the error bars to the SD. D. Histological characterization on frozen tissues of mice deposits with CR staining revealed the birefringence characteristic of amyloid deposits (top) and their fluorescence (bottom) of the CR and anti hλ-LCs antibody in the heart, the spleen and in the kidneys. Example of a mouse analyzed at 5 months after induction (high score).

### Long-term follow up revealed exceptional spontaneous AL amyloidosis development with age

In humans, AL amyloidosis can take several years to develop (*49*). We also showed that even when seeded with AL fibrils or amyloidogenic VLs, detectable amyloid deposits often occur after several months in the mice, corroborating findings in other mouse models of systemic amyloidosis (*34, 47*). Consequently, we conducted a long-term follow-up on 28 λS-DH mice. Since we never detected any sign of morbidity and we did not find at this time any biomarker that could help to detect the onset of the disease, we randomly sacrificed mice at 12-16 months (n=9), 18-24 months (n=14) and > 24 months (n=5). 2 out of 28 mice developed AL amyloidosis at the age of 14 and 18 months (**Supp Fig. S5**). This indicates that spontaneous amyloidosis may rarely happen in these mice but the rarity of this event does not allow us to conclude that amyloidosis develops with age or that it is linked to any precursor event.

### Deep characterization of amyloid deposits shows striking similarities with those from AL patients

In order to further analyze the composition of the amyloid deposits in our mice, we performed fibrils purification from the cardiac tissue of positive mice using the “Pras” protocol as previously published (*50*). The presence of fibrils was confirmed by atomic force microscopy (AFM) (**Fig. 6A**). The main constituent of the fibrils was, as expected, the λS-LC, and more precisely a fragment recognized by the anti-IGLV6 antibody (**Fig. 6B**). However, the size of the main VL fragment was shorter than the rλS-VL. As expected, mass spectrometry analysis of purified fibrils readily detected the λS-LC, with peptides both from the VL and from the CL parts of the protein, together with common amyloid-associated proteins, such as ApoA4, ApoE and vitronectin (*51*) (**Figure 6C**). However, we did not detect Serum Amyloid P component (SAP). We also detected Collagen VI subunits that have been recently described to be structurally associated to amyloid fibrils in the heart of an AL amyloidosis patient (*52*). Some studies have shown that, during fibrils extractions by the Pras protocol, tryptic activity of collagenase, used to dissociate the tissues, can degrade parts of the LCs that are not involved in the interactions within the core of the fibrils (*15*). Consequently, to better characterized the composition of the amyloid fibrils in AL positive hearts, we also used a gentle extraction procedure in presence of a protease inhibitor cocktail that leads to an enrichment of unsoluble content of the tissue as previously described (*15*). Amyloid deposits were enriched from three soluble VL-induced λS-DH AL positive mice. As a control, two amyloid-negative samples (one non-induced λS-DH mouse and one non-transgenic DH) were subjected to the same extraction procedure. Immunoblotting showed no immunoreactive bands in the extracts from amyloid-negative mice, while multiple ones were visible in the amyloid-positive animals, spanning from approximately 25 to 12-15 kDa and consistent with the presence of the full-length light chain and fragments (**Fig. S6A**). This confirms that the endogenous λS-LCs from mice can elongate the λS-VL seeds preformed either *in vitro* or *in vivo.* The amyloid-positive samples were then analyzed by 2D-PAGE and 2D western blotting to resolve the light chain proteoforms (**Fig. 6D**). The fragmentation pattern of the λS-LC in fibrils was strikingly similar in all the mice tested, but also to the one previously observed in the heart of a patient with AL amyloidosis and sharing the same IGLV6-57-derived LC (AL55) (**Fig. S6B**) (*15, 16*). The spots on the Coomassie-stained gels matching the western blot signals were excised, trypsin-digested and analyzed by LC-MS/MS (**Fig. 6E**). Mass spectrometry showed that the 25 kDa spots contain tryptic peptides from the entire LC sequence, except for the C-terminal one, which was not detectable. The lower MW spots, instead, contain the entire variable region and progressively shorter portions of the constant domain. Some spots with MW <12-15 kDa, which are not detectable by WB, were shown to contain exclusively the VL or portions of it (**Fig. 6E**). Taken together, our results confirm that the VL is the major common component of the AL amyloid deposits but that endogenous full length λS-LC participates in the formation of the insoluble material extracted from the tissue. The similarities in the fragmentation pattern of the LCs between patients and our mouse model also suggest a conserved mechanism of amyloid fibrils formation.

**Figure 6:**
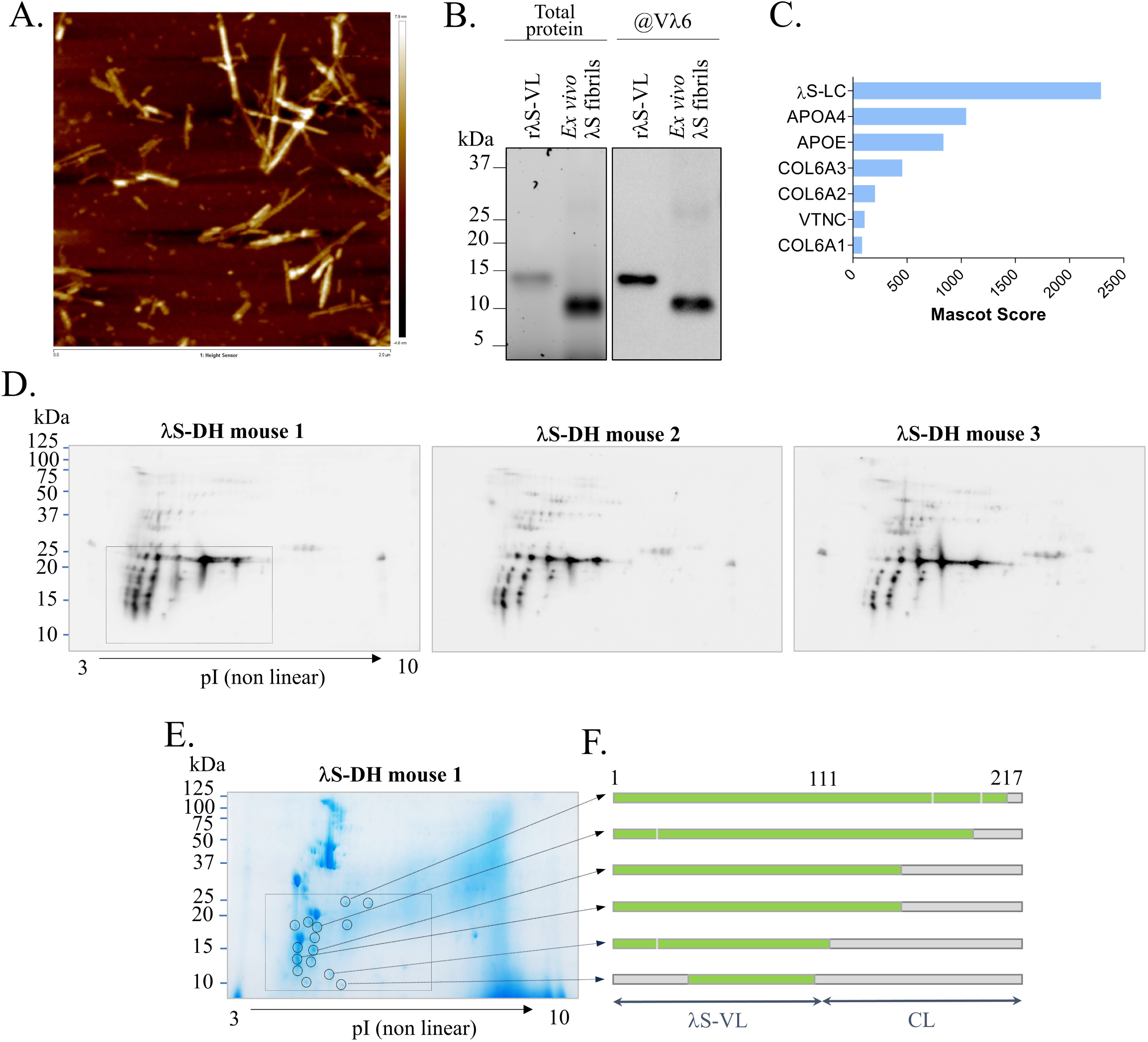
Molecular characteristics of the amyloid deposits in λS-DH mice. A. The presence of fibrils in the samples upon extraction and enrichment from the cardiac tissue of mice was confirmed by AFM. The fractions containing the highest concentrations of fibrils were pooled and used for further experiments. The image frame is 2 * 2 µm. The height sensor is 12.5nm (−4.6 to 7.9 nm). B. Analysis of the purified λS-DH *ex-vivo* fibrils by SDS-Page and Western Blot. Total protein was revealed by the Stain-Free technology (Biorad). Blotting was done with an anti-human Vλ6-57 antibody. C. Samples from (A) were analyzed by mass spectrometry. The protein content of samples was given by the Mascot score. Only the proteins associated to the amyloid deposits are shown in the representation. D. Analysis of deposited light chains in amyloid-positive mice by 2D-PAGE and 2D western blotting using anti-hλ LCs showed a complex population of immunoreactive trains of spots in all mice (25 to 12 kDa). The appearance of these spots was almost superimposable across the animals. The boxed region in mouse 1 matches the corresponding region shown in (E). E. Coomassie-stained gel from λS-DH mouse 1. The spots whose position matches the immunoreactive spots in the western blot shown in (A) (circled) were excised, trypsin-digested and analyzed by LC-MS/MS. F. Coverage of the monoclonal light chain sequence in each spot, from the tryptic peptides identified by MS.

### AL fibrils deposition induces early cardiac dysfunction in mice

Having shown that λS-DH mice develop AL amyloidosis in heart upon induction, we sought to determine if these deposits could lead to cardiac dysfunction. Cardiac amyloidosis diagnosis in patients is based mainly in the detection of cardiac failure biomarkers in the bloodstream, the more sensitive and specific being NT-proBNP, or echocardiography. Unfortunately, at the time most experiments of the present study were done, we could not find a reliable NT-proBNP assay kit for mouse. Those tested showed high assay variability in control groups and inconsistencies between kits, ranging from pg to ng on the same samples. Recently, we eventually found a new assay, which gave more reliable results and we retrospectively measured the concentration of NT-proBNP in the serum of mice with confirmed cardiac amyloidosis (AL, 2.80, range=2-3, n=10) (**Fig. S7A**) and compared it to the levels in DH mice (DH, n=8) and λS-DH mice without deposits (LS, n=8). We observed a significant increase between the AL positive λS-DH and the AL-mice (mean 1633 pg/ml +/- 400 vs 237 pg/ml +/- 42 respectively, p=0.0009), and between the λS-DH AL+ and the DH mice (332 +/- 73.9, p=0.0085) (**Fig. 7A**) albeit with a high heterogeneity in the AL positive group. No significant difference was observed between the λS-DH without deposits and the DH mice (p=0.328).

**Figure 7:**
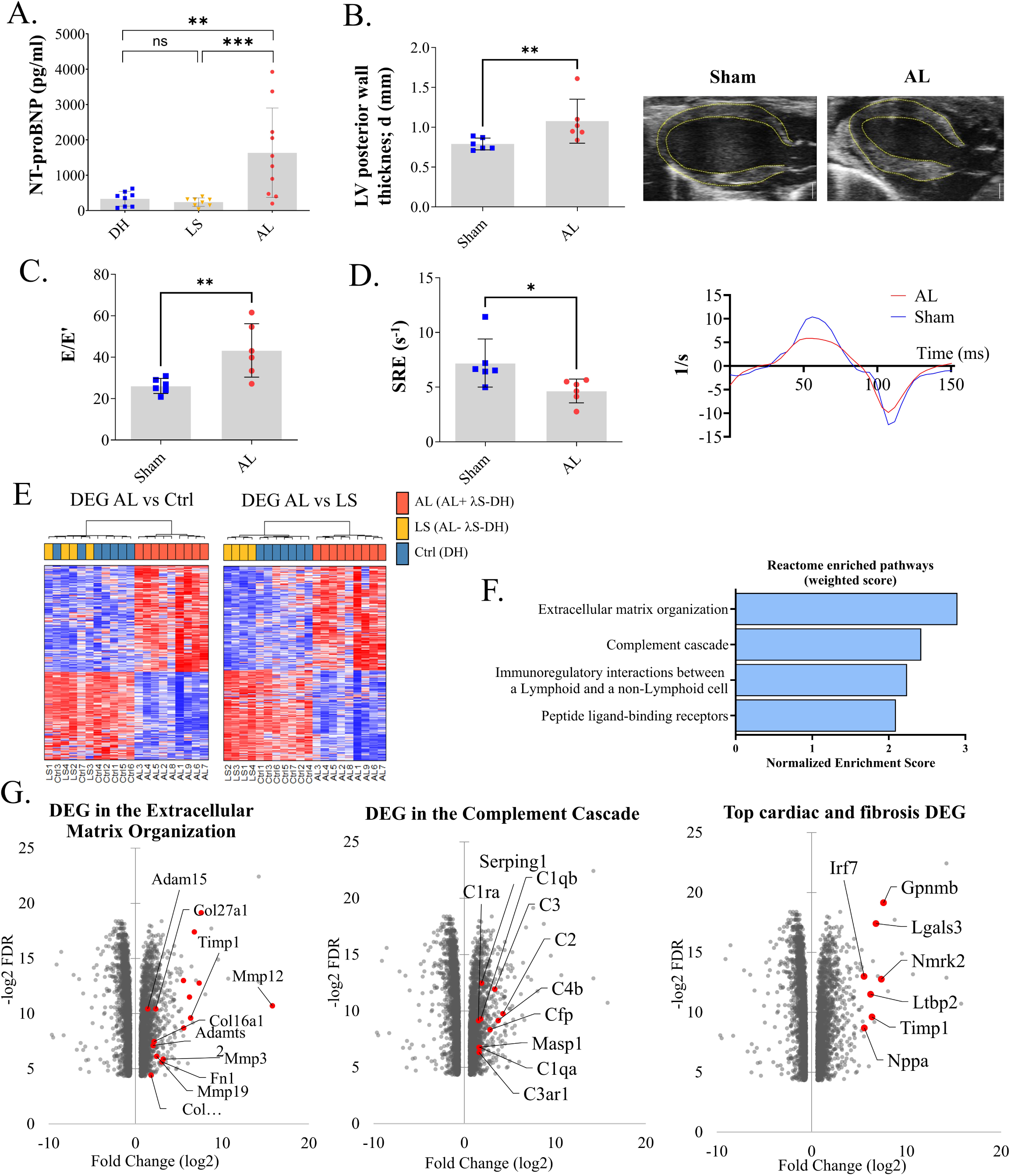
Analysis of the cardiac function related to amyloid deposition in λS-DH mice. A. Plasma NT-proBNP was dosed in λS-DH with a mid-high score of cardiac amyloid deposition (AL+, n=10) as well as controls, λS-DH mice without amyloid deposition (AL-, n=8) and with DH mice (n=8). Significant 7-fold and 5-fold increases were observed when comparing the λS-DH AL+ mice with the AL-λS-DH (p=0.0009) and DH (p=0.0085) mice respectively. No significant difference was observed between AL-λS-DH and DH mice (p=0.328). Bar graphs correspond to the mean and the error bars to SD. B. Cardiac measurement of the diastolic (d) left ventricular (LV) posterior wall by Ultrasound analysis in age-matched DH control mice (Sham, n=6) and λS-DH with a high score of cardiac amyloid deposition (AL+, n=6) showing the increase of the ventricular wall thickness (p= 0.0087) (right). Representative image of the long axis LV segmentation (left). C. Cardiac filling pressure, given by the E/e’ ratio, was significantly increased in the AL+ cohort (p=0.0087). D. Cardiac early diastolic strain rate (SRE) and representative curves in AL+ and control mice showed a significant decrease of the ventricular relaxation time correlated to the amyloid deposits (p = 0.0152). E. Heatmaps showing the differentially expressed genes (DEG, fold change, FC < |1.5| and False Discovery Rate, FDR < 0.05) among the 10059 genes analyzed in λS-DH mice with cardiac deposits (AL) compared to DH mice (Ctrl, left) and compared to λS-DH mice without deposits (LS, right). Mice were clustered according to the DE profiles. Red corresponds to overexpressed genes and blue to downregulated genes. F. Reactome enriched pathways of the 3887 DEG from this analysis. The represented pathways are the result of the weighted score enrichment of significantly enriched pathways (n=13) (FDR < 0.05) G. Some of the overexpressed genes associated to the extracellular matrix organization and complement cascade pathways are annotated in red on the volcanoplot according to their Fold change (log2) and their adjusted p-value (represented by the −log2FDR). 7 genes in the Top 20 of overexpressed genes in our analysis were associated with cardiac dysfunction or correlated to the fibrotic processes.

We also analyzed by high-resolution echocardiography two cohorts of 4 λS-DH mice 6 months after VL-seeding induction (12-18 month-old at sacrifice) and 8 age-matched controls (DH). Then, mice were sacrificed for histological studies. Upon the AL mice, one did not present AL deposits at sacrifice and another showed evidence of an intracardiac thrombosis (**Fig. S7B**). Both were excluded from the echocardiography analysis. Similarly, in the control group, one died for undetermined reason before the end of the experiment and another one presented with a significant tumor mass in the liver and a right kidney atrophy. Both were excluded from the analysis. As expected from previous experiments, we observed a high heterogeneity in the amyloid burden of positive AL mice (mean score 2.5, range=1-3) **(Fig S7C)**. We analyzed classical parameters used in human to characterize cardiac AL amyloidosis (*53*). Despite the heterogeneity of the amyloid burden, echocardiographic parameters were consistent with ventricular hypertrophy with an increased diastolic posterior wall thickness (LVPWd) in AL positive hearts (1.08 mm +/- 0.112 vs 0.790 +/- 0.030 in controls, p = 0.0087) (**Fig. 7B)** and a tendency that did not reached significance for global ventricular hypertrophy (corrected LV mass, p = 0.0649) (**Fig. S7D-E**). We also detected an early diastolic dysfunction in AL positive mice, including higher left ventricular (LV) filling pressures (E/e’) (43.3 +/- 5.26 vs 26.1+/- 1.48 in Controls, p = 0.0087) (**Fig. 7C**) and lower early diastolic strain rate (SRe) (4.658 +/- 0.443 vs 7.207 +/- 0.897, p = 0.0152), suggesting a reduction in passive ventricular relaxation (**Fig. 7D**) associated with a preserved ventricular ejection fraction (58.6% +/- 5.56 vs 56.8% +/- 1.72, p = 0.8182) (**Fig. S7F**) as expected in AL amyloidosis. However, the global longitudinal strain (GLS) and early to late diastolic transmitral flow velocity (E/A) were non-significant between groups (p=0.7879 and p = 0.6753, respectively) (**Fig. S7G**). A reduced ventricular filling capacity was observed by the increase in left atrial (LA) volume and a decrease in LA ejection fraction (**Fig. S7H**). All these data are mainly indicators of heart failure with preserved ejection fraction. Interestingly, as indicated above, one AL-induced mouse presented with an intracardiac thrombus in the left atrium (**Fig. S7B**) which is a frequent observation in patients suffering with cardiac AL amyloidosis (*54*). Histological studies confirmed a high CR positive amyloid burden in the cardiac tissue of this mouse.

### Transcriptomic analyses define extracellular molecular events but not cellular toxicity as a feature of amyloid fibrils toxicity in mice

To further decipher the molecular mechanisms leading to cardiac dysfunction and toxicity, we performed a transcriptomic analysis on heart apical tissue from the λS-DH transgenic mice using bulk RNA sequencing. We analyzed cardiac tissue from AL positive mice (n = 9, AL1 to 9, mean amyloid score 2.33, range=1-3 (**Fig. S8A**) and age-matched DH mice as controls (n = 7, Ctrl 1 to 7). Since it has been shown that a soluble full-length amyloidogenic LC could exert a direct toxicity on cardiac cells (*21, 55, 22, 56*), we also analyzed non–induced amyloid negative cardiac tissue from λS-DH (n = 4, LS 1 to 4). For differentially expressed genes (DEG), we applied a minimum fold change of 1.5 (Log2FC = 0.58) and a false discovery rate (FDR) < 0.05. Principal Component Analysis (PCA) and hierarchical clustering of the samples showed a clear separation between AL mice and the other groups (LS and Ctrl) (**Fig. 7E** and **Fig. S8B**). However, the two AL negative groups showed no clear distinction, whatever the chosen comparison (AL vs Ctrl or AL vs LS). The transcriptional changes (DEG) between these three groups revealed that, among 10059 significantly expressed genes, 1135 (11.3%) were differentially expressed between AL and Ctrl samples and 1766 (17.6%) between AL and non-amyloid LS λS-DH mice. Comparison between LS and Ctrl confirmed the PCA and hierarchical clustering with only 3 significantly deregulated (0.03%) genes with irrelevant functions (**Table S1**). Since LS and Ctrl groups were not distinguishable, we decided to merge them in a unique group of control samples to strengthen our statistical analysis. This analysis resulted in 3887 genes (39%) which were differentially expressed (1893 upregulated and 1994 downregulated genes) (**Fig. S8C**). We used WEB-based Gene Set Analysis Toolkit (WebGestalt) (*57*) to study the deregulated pathways (Reactome database) (*58*) and gene ontology (GO) Molecular Function (MF), Cellular Component (CC) and Biological Process (BP) signatures of the amyloidogenic cardiac tissue. Despite the number of DEG, only 13 pathways, all positively enriched, were significant (FDR < 0.05) (**Fig. S8D**) and many of them were closely related so that we used the weighted set cover option to avoid redundancy (**Fig. 7F**). The most enriched pathways were related the extracellular matrix (EM) organization (7 out of 13 statistically enriched pathways), with 55 differentially expressed genes encoding for endoproteases (MMPs, ADAMs and extracellular cathepsins) and their inhibitors (TIMPs), as well as EM-related proteins, such as collagens, fibronectin, laminin and adhesion molecules (**Fig. 7G and S8F**). Other pathways comprised complement cascade (4 out of 13 enriched pathways), Immunoregulatory interactions between a Lymphoid and a non-Lymphoid cell and Peptide ligand-binding receptors (**Fig. 7F**). The analysis of the Gene Ontology processes associated to the DEG corroborated these findings, and innate immune cell populations were principally enriched in the cell-type landmarks analyzed by the Mouse Cell Atlas (**Fig. S8G**). Strikingly, we did not identify any obvious molecular signatures of cellular toxicity in the amyloid cardiac tissue. However, we detected several known markers of cardiac dysfunction and cardiac fibrosis among the top 20 overexpressed genes (**Fig. 7G**) including *Nppa, Lgals3, Gpnmb, Timp1, Nmrk2, Ltbp2, Irf7* (*59–63*). Most of these markers were also overexpressed in AL positive hearts compared to each control group (LS or Ctrl) independently (**Table 2**). It is noteworthy that while *Nppa* expression was found significantly correlated to amyloid deposits in our mice (Log2 Fold Change = 5.57 and False Discovery Ratio = 0.002), *Nppb* was not significantly overexpressed (Log2FC = 2.18 and FDR = 0.054), likely due to the high variability of its expression between mice. Collectively, these findings indicate that the soluble amyloidogenic λS-LC does not seem to exert any obvious toxic effect on the cardiac tissue of our transgenic mice. Nevertheless, amyloid deposition demonstrated clear effects on the extracellular compartment of cardiac tissue, consistent with early fibrotic processes and immune cell infiltration.

## DISCUSSION

We herein describe the first transgenic mouse model of systemic AL amyloidosis with dominant cardiac involvement. To achieve a pathogenic production level of the amyloid LC, the main limitation of previous models (*33, 31, 32*), we applied a strategy already successful in another monoclonal LC-associated disease (*39*), coupling an orthotropic production of the human LC by plasma cells and the removal of endogenous HC to force the production of free LCs. These mice continuously produce a level of fLC that is above the median level observed in AL patients (*64*) and approximatively four fold more than in the patient from whom the LC gene was isolated. With rare unexplained exceptions, mice did not develop spontaneous amyloidosis but upon induction, they accumulated AL amyloid fibrils in heart, spleen and, to a lower extent, in other visceral organs. These fibrils have typical pathological features of λ LC AL amyloid deposits with most of the classical amyloid signature proteins (ApoE, ApoA-IV, Vitronectin). To achieve amyloid deposition in λS-DH mice, we used two different induction methods. One is the classical seeding strategy with injection of preformed fibrils that accelerates amyloid formation by bypassing the long early stages of nucleation. In our case, and contrary to most other models, which use purified *ex vivo* fibrils, we injected *in vitro* fibrils exclusively composed of the recombinant VL part of λS-LC. We used this approach for two reasons: the unavailability of tissue from λS patient to make *ex vivo* fibrils and our failure to obtain any fibrils from the recombinant full length λS-LC. For the second protocol, we injected the soluble VL alone assuming that its intrinsic amyloidoigenicity would allow its aggregation *in vivo*. Both protocols led to amyloid deposits in mice, but with different timing and incomplete penetrance. Since the fibrils injections were able to trigger a rapid burst of amyloidosis, detectable as soon as 48h after induction, the VL injections led to a slower, more physiological development of the disease. Due to the rapid renal clearance of low molecular weight proteins, we assume that few oligomers of the VL are quickly formed in the circulation or tissues where they remain to be progressively elongated by full length LC and spread to the other organs through secondary nucleation and fragmentation (*65*). This means that once formed, these oligomers quickly become resistant to proteolysis. This is, to our knowledge, the first *in vivo* evidence of the enhanced potential of the VL to initiate the subsequent formation of mature amyloid fibrils elongated by full length LC. Of course, we conveniently used the strict VL fragment for our experiments based on our *in vitro* fibrilization tests, the previously cited *in vitro* data and the recent information obtained from 3D structures of fibrils from AL patients, which showed that the core of AL fibrils are overall established by the VLs (*14, 66, 67*). However, we assume that the pathophysiological process in human is likely more complex than a simple, precise, release of the sole VL part. Accordingly, Lavatelli *et al.* recently showed *in vitro* that a partial degradation of the constant domain, as frequently observed in AL deposits, is sufficient to unleash the amyloidogenic propensities of an otherwise stable full length LC. Consequently, based on our findings, it is tempting to argue that a partial proteolysis of fLCs, releasing fragments containing destabilized VL that can rapidly form the first nuclei, could occur in serum or extracellular matrix of organs and could be required to initiate the elongation of amyloid fibrils with the circulating monoclonal full-length LC. With this in mind, it would be interesting to identify the potential molecular players of this LC fragmentation, their location and activation.

With very rare exceptions, mice do not spontaneously develop AL amyloidosis, as previously shown in other systemic amyloidosis models (*47*). There are several hypotheses to explain this observation, including a faster turnover rate of proteins or a better control of proteostasis in mice (*35, 36*). However, in human, the production of fLC precedes from several years the occurrence of symptomatic AL amyloidosis (*49*) and in other systemic amyloidosis, especially hereditary ones, the onset of the disease appear usually after several decades. Accordingly, in λS-DH mice, amyloidosis development takes several months even when accelerated with a large quantity of amyloidogenic VL. Then, we cannot exclude that even with the pathogenic level of fLC observed in human, mice simply do not live long enough to let the time to allow overt amyloid deposition to develop. Since an increase of fLC level precedes the onset of clinical symptoms, reaching higher levels of fLCs in future transgenic mice could help accelerate the natural onset of the disease.

Characterization of mature amyloid fibrils in tissues or extracted from hearts of positive λS-DH mice allowed some important considerations regarding the composition of cardiac amyloid and its comparison with its human counterpart. In first place, these analyses showed that the insoluble deposits do not exclusively contain the VL used for seeding, but they incontrovertibly indicate that the endogenous full-length LC is also incorporated. This observation opens fundamental perspectives on the mechanisms of amyloid formation *in vivo*. In particular, it indicates that a critical concentration of VL-containing fragments is needed for priming the process, but that the full-length LC subsequently or concomitantly participates to amyloid progression. Although the events that generate the complex population of deposited LC fragments are not known, these proteolytic events are virtually identical in the different animals. Moreover, the biochemical similarity between mice and human LC deposits is remarkable. As in human amyloid, major fragments in mice contain the complete VL and progressively shorter stretches of the CL, suggesting that the VL constitutes the core of the murine AL amyloid fibrils, as previously shown in human (*66, 68*). The pattern also suggests that the proteolytic remodeling mechanisms associated to LC aggregation and/or to aggregate metabolism are analogous in mice and human. The transgenic λS-DH mouse is therefore a representative model and a useful resource to study the formation, composition and metabolism of human AL deposits.

In contrast, the organ tropism seems to be relatively distinct in the mice compared to the patient from whom the transgenic LC was extracted. Amyloidosis in λS-DH mice develops mainly in heart and associated vessels. At odds with the patient who was first diagnosed with renal involvement, the deposits in kidneys appear late and remain restricted to a few isolated glomeruli. Although the patient presented with typical cardiac dysfunction at the time of diagnosis, understanding this discrepancy in organ tropism between human and mouse is likely to provide clues about the mechanisms leading to a preferential organ involvement.

In addition to the formation of amyloid deposits, this model also develop early symptoms of cardiac dysfunction. Due to the inconstant penetrance of the disease, the variability of deposits progression and possibly the heterogeneity of the genetic background, it remains difficult to associate the amyloid burden to the severity of the cardiac dysfunction. However, we found a significant increase of NT-proBNP, the most reliable plasmatic biomarker for AL amyloidosis, in AL positive mice before sacrifice compared to non-induced λS-DH or DH control mice. Induced λS-DH developed also early signs of diastolic dysfunction with preserved ejection fraction. Although incomplete, this dysfunction shows that beyond AL deposits, λS-DH can also be a relevant model to study the molecular events leading to cardiac toxicity. Accordingly, transcriptomic analyses showed significant changes in several genes classically overexpressed during cardiac dysfunction, particularly those involved in fibrosis and extracellular matrix remodeling (*63*). Among these genes, it is interesting to note the presence of many metalloproteinases and extracellular cathepsins that could also participate in the proteolytic events leading to LC fragmentation. Although we did not detected significant infiltration of immune cells in the cardiac tissue, it seems that myeloid-derived cells, including macrophages, monocytes or dendritic cells, together with cardiac fibroblasts may account for this profibrotic phenotype. Deciphering the origin (cell type), location and function of these proteases will likely give clues about their role during AL amyloidosis development. For this purpose, spatial transcriptomic and/or proteomic on mouse tissues with different degrees of amyloid deposition would likely be of great value, especially to better understand changes in cell populations and their activity in the vicinity of amyloid deposits.

There are two major discrepancies with previous *in vitro* or non-mammalian models. First, based on transcriptomic analysis and histology, there is no clear indication of cardiomyocytes toxicity, induced by the fibrils. We may easily hypothesize that mice only develop the early stages of the disease. Longer exposure and further accumulation of deposits would ultimately lead to myocyte damage. Accordingly, AL positive mice showed only discreet cardiac hypertrophy and the amyloid load in mice hearts, even with the highest scores, remains below that observed in human cardiac biopsies (*69*). The second discrepancy concerns the apparent absence of cardiac toxicity caused by the soluble amyloidogenic LC. The toxicity of amyloidogenic fLCs has been extensively studied on cardiac cells (cardiomyocytes, cardiac fibroblasts and mesenchymal cells) (*70, 71, 22, 72, 73*) *in vitro*, as well as and in lower animal models, such as *C. elegans* and zebrafish. Among the observed effects, mitochondrial dysfunction, ROS production and cell death were directly associated with soluble fLCs. In the present mouse model, the production of the LC by plasma cells continuously exposes cardiac cells to a high level of an amyloidogenic LC. However, mice with no amyloid deposits seem healthy, with no increase of NT-proBNP and a transcriptional landscape of cardiac tissue almost identical to control mice. There are several hypotheses that may be proposed to explain this difference. First, the patient from whom the λS-LC gene was extracted was firstly diagnosed with renal involvement so one could hypothesize that it is not a cardiotoxic LC *per se*. However, even in the absence of direct evidence of cardiac deposits in the patient, his clinical symptoms were characteristic of AL amyloidosis-induced cardiac dysfunction. Then obviously, mice are not men, and differences in protein turnover, extracellular protein chaperoning, cellular resistance to LC-induced stress or absence of LC binding to cell surface could all account for the undetectable LC toxicity in our mouse model (*22, 35, 36, 73, 74*). On the other hand, the direct toxicity of amyloid LCs on cultured cells, *ex vivo* organs or in lower animals could also be challenged since these systems do not reproduce the *in vivo* extracellular proteostasis, including circulating chaperones and proteases, which could protect cells from unstable or unfolded dangerous LCs in the mice or in humans (*75*). In addition, immune system, extracellular matrix environment or cell type diversity, all potentially involved both in the pathogenicity but also the protection of organs from toxic LCs, cannot be reproduce *in vitro* or in lower organisms. Finally, even in absence of direct toxicity, λS-DH mice develop an early cardiac dysfunction closely resembling the human disease. This dysfunction relies on the presence and accumulation of amyloid fibrils that progressively replace and remodel the extracellular space of cardiac muscle, likely causing mechanical constraints as a first step. Then, we can hypothesize that further accumulation of amyloid fibrils, reaching a critical threshold, would ultimately lead to cellular toxicity and more severe cardiac dysfunction. The quick recovery of the cardiac function in patients upon the reduction of circulating LC remains to be understood but could rely on the arrest of further accumulation of new toxic fibrils.

In addition to its contribution to the pathophysiological mechanisms of AL amyloidosis, our model also fills a crucial gap in the validation of new therapeutic approaches. Having shown that amyloid deposits are very similar to those observed in humans, both in terms of localization and composition, it is now possible to use this model to test new therapeutic approaches designed to eliminate amyloid fibrils in tissues, or to stabilize / inhibit the circulating amyloid precursor. The absence of Ig heavy chain, and therefore of a proper humoral immune response, is of particular interest in this respect, as it allows the repeated injection of humanized molecules without any risk of immune rejection. However, the main limitation to the use of this model for pre-clinical purposes remains the incomplete penetrance of the disease. Adapting the induction protocol by repeated seeding or *ex vivo* injection of fibrils obtained from previous positive mice is currently being evaluated and we are confident that this will rapidly work.

We describe here the first mouse model recapitulating most of the features of a systemic AL amyloidosis with cardiac involvement, including progressive accumulation of fibrils in the extracellular space of organs and early typical cardiac dysfunction. We confirm the crucial role of the variable domain in initiating amyloid deposits and provide new insights into the toxicity of amyloid fibrils for heart. This model offers a new avenue for research on AL amyloidosis and fills an important gap for the development of new therapies.

## MATERIALS AND METHODS

### LC gene extraction and sequencing

The human monoclonal LC genes used in the study were obtained from bone marrow (BM) aspirates as previously described (*76, 77*). The AL λLC cDNA was extracted from a patient with biopsy-proven AL amyloidosis (λS-PT). The LC is derived from the Vλ6-57 germline gene (95,53% homologous) rearranged on a Jλ3*02 junction segment (92,11% homologous) and a constant Cλ3 domain. The κR-LC gene used to generate the κR-DH model was extracted from a patient with multiple myeloma and cast nephropathy, with no amyloidosis in the kidney biopsy and no other clinical argument to suspect amyloidosis.

### Transgenic mice models generation

To generate the transgenic mice models (λS-DH and κR-DH) two similar strategies were used as previously described (*38, 77*). For λS-DH, the complete cDNA coding for the selected human monoclonal λLC was introduced in place of the mouse Jκ segments in the κ locus thus generating a fully human LC. For κR-DH, only the cDNA coding for the variable/junction domains (VJ) was introduced, generating in the mouse a chimeric human/mouse LC composed of the human VJ domain and the mouse Cκ domain as previously described (*38, 77*).

Mice were crossed with DH-LMP2A mice, kindly provided by S. Casola (IFOM, Milan, Italy) (*42*). The transgenic strategy for λS-DH is showed in **Figure 1A** and was previously described (*78*). The two models (λS-DH and κR-DH) produced similar levels of free LC. Mice were maintained in pathogen-free conditions, with food and water *ad libitum*, unless otherwise stated. All experimental procedures have been approved by our institutional review board for animal experimentation and of the French Ministry of Research (APAFIS #7655-2016112211028184).

### LC dosing

Serum were analyzed for the presence of FLC using Freelite^TM^ (The Binding Site, Birmingham, UK) assay on BNII nephelometer (Siemens healthcare, Herlangen, Germany) to the appropriate dilution for each sample and according to according to the manufacturer’s instructions.

### Western Blot

Proteins were separated by reducing or non-reducing SDS-PAGE on Mini-Protean TGX Stain-free gels (Biorad). Total protein were detected by the fluorescence of the Stain-Free technology. After the proteins were transferred onto polyvinylidene difluoride membranes (Millipore), membranes were blocked in 5% milk Trisbuffered saline (TBS), following by incubation with desired antibodies in 3% milk TBS (**Table 2**), washed three times with TBS 0.1 % Tween and revealed by chemiluminescence (ECL, Pierce).

### Flow cytometry

Intracellular staining was performed using the Intraprep™ kit (Beckman Coulter). Flow cytometry analysis were performed on a CytoFLEX (Beckman Coulter) and data were analyzed with Kaluza Software (Beckman Coulter). Corresponding antibodies are listed in the **Table 1**.

### Histological studies

Organ samples were processed for immunofluorescence (IF) studies, as previously described (*76*). In brief, IF was performed on OCT-included organs and snap frozen in isopentane using a Snap Frost 2 (Excilone). Cryosections of 9µm were fixed with cold acetone, blocked with phosphate-buffered saline (PBS) and bovine serum albumin (BSA) 3% and then stained with appropriate antibodies (**Table 1**). For Congo Red (CR) staining, 2.87mM CR alkaline solution was freshly prepared by adding NaOH 1% following a five-minute staining before washing with PBS. Slides were observed on a NiE microscope (Nikon). For electron microscopy (EM) and immunogold studies, tissues were fixed in 3% glutaraldehyde in 0,1M phosphate buffer (pH 7,2) at 4°C and processed as previously described (*76, 77*). For histochemical staining, tissues were fixed in 4% paraformaldehyde and paraffin-embedded as previously described (*77*).

### Score of amyloid deposition in mice

The score was estimated by the fluorescence of the Congo Red in the hearts of mice as follows: score low corresponds to one or more focal deposits in the myocardium and/or blood vessels; mid corresponds to some diffuse deposits within the myocardium; and high corresponds to diffuse deposits throughout the myocardium.

### Biochemical parameters determination

Biochemical parameters were measured on overnight urine collection and plasma samples, obtained by retro-orbital puncture under anesthesia or by cardiac puncture after their euthanasia on heparin. Plasma concentrations of creatinine were measured on a Konelab 30 analyser with a creatinine enzymatic test (ThermoFisher Scientific). Urine albumin concentrations were measured using an albumin mouse ELISA kit (Abcam), according to the manufacturer’s recommendations. Mouse N-Terminal Pro-Brain Natriuretic Peptide (NT-proBNP) was estimated in plasma using a mouse NT-proBNP ELISA kit (Elabscience).

### Cell line generation

To express the λS-LC *in vitro*, the corresponding cDNA obtained from the patient was amplified and cloned in a modified pCpGfree plasmid (Invivogen, San Diego, USA) containing the neomycin resistance gene. Series of transfections in the murine hybridoma/myeloma cell line SP2/0 (ATCC® CRL-1581™) were performed using Cell line Nucleofactor kit V (Amaxa/Lonza, Basel, Switzerland) and a Nucleofactor II device (Amaxa/Lonza, Basel, Switzerland). Positive clones were then selected using neomycin (1mg/mL, Fisher Bioreagent, Pittsburg, USA).

### Expression and purification of human recombinant **λ**S Full-Length free LC

Full-length free LC secretion in 7-day culture supernatant was quantified as previously described using ELISA. The selected best producing SP2/0 λS-LC clone was cultured in a mini-bioreactor system (CELLine CL 350, Integra Biosciences) and the secreted FLC collected according to manufacturer’s instructions. All collected samples were pooled and FLC was purified by Binding Site (Birmingham, UK) or by affinity chromatography using a LambdaFabSelect column (Cytiva) or a polyclonal anti-human lambda (Dako) antibody-containing resin. Protein purity was assessed by SDS-PAGE and Coomassie Blue staining. The resulting purified protein concentration was estimated as previously described in the “LC dosing”.

### Expression and purification of recombinant human Variable Domain (r**λ**S-VL)

Recombinant human λS-VL and was periplasmatically expressed in *E.coli* BL21 competent cells based on the pET12a vector (Novagene) containing an optimal peptide for periplasmic expression. After overnight growth, the proteins were extracted with a cold osmotic shock and purified on a POROS™ XQ anion exchange resin (Thermo Fisher Scientific) followed by a Ceramic Hydroxyapatite resin on an AKTA-FPLC system (GE Healthcare). Protein purity was assessed by SDS-PAGE and Coomassie Blue staining and the concentration was determined by BCA assay (Thermo Fisher Scientific). Samples were stored in 10 mM Hepes, 100 mM NaCl, pH 7.6.

### Size Exclusion Chromatography on r**λ**S-VL

The native SEC-UV/MS was carried out using an ACQUITY UPLC Protein BEH200 SEC column (4.6 x 150 mm, 1.7 mm particle size; Waters, Milford, MA, USA). An isocratic elution with water at 0.1 mL/min was used for chromatographic separation on a Nexera LC40 system (Shimadzu, Noisiel, France) equipped with UV detection at 280 nm. Sample injection amounts of 10µg were used and data acquisition was controlled by Hystar (Bruker Daltonics).

### Heparin induced fibrilization of r**λ**S-VL or r**λ**S-LC

For aggregation assays, 33 µM of protein (λS-VL or LC) was incubated in aggregation buffer (10 mM Hepes, 100 mM NaCl, 5 mM DTT, 0.8 µg/µl Heparin, pH 7.6) at 37°C for the length of the experiment at 37°C and 300 rpm. For the *in vitro* seeding assays, rλS-VL seeds were prepared by sonication of λS-VL fibrils 20 seconds at an amplitude of 90% using an Ultrasonic homogenizer (Bandelin). Addition of 1.6 µM seeds to the λS-VL aggregation assay mentioned above was done at the beginning of the reaction. 1.6 µM λS-VL fibril seeds prepared as mentioned above were incubated with 6.5 µM rλS-LC in aggregation buffer for 4 days at 37°C, 300 rpm. Fibrillation kinetics was followed by ThT fluorescence (FLUOstar Omega, BMG Biotech), and confirmed by Atomic Force Microscopy (AFM) or Transmission Electron Microscopy (TEM).

### Visualization of amyloid fibrils

For TEM analysis, 10 μl of fibril sample was deposited on a formvar/carbon grid of 200 copper mesh (Agar Scientific), and excess liquid was removed. Negative staining was performed with Uranyless 30% ethanol (Em-Grade), washed twice with H2O, and air-dried. Grids were analyzed on a JEM-1400 Flash Transmission Electron Microscope. For AFM analysis, 10 µl of a fibril sample was applied to a freshly cleaved mica surface (Plano GmbH). After 3 min incubation, the mica was washed with water and dried under air flow. The sample was scanned using the tapping mode on the AFM 5500LS (Agilent).

### Thermal stability of the proteins r**λ**S-VL or r**λ**S-LC

The concentration of the proteins was adjusted at 100 µg/ml in Hepes buffer (10 mM Hepes, 100 mM NaCl, pH 7.6) and the thermal stability was measured on a Tycho NT.6 (NanoTemper Technologies) between 35 and 95 °C. The thermal unfolding midpoints (T_m_) were calculated using the first derivative of the fluorescence intensity ratio at 350 nm /330 nm, corresponding to a conformational shift of tryptophan residues.

### Induction of the amyloid deposition in mice

λS-DH mice between 3 and 12 months old were injected by i.v. with 200µl of 2 mg/ml rλS-VL seeds (sonicated fibrils) or 4 mg/ml soluble rλS-VL in a PBS solution. Mice were euthanized at different timepoints, going from 48h after injection to 12 months. Organs samples were immediately frozen in −80°C isopentane embedded or not in Tissue-Tek O.C.T (Sakura), or fixed in 4% paraformaldehyde and paraffine-embedded for histological studies. As controls, κR-DH (human monoclonal FLCs) mice, DH-LMP2A (mouse polyclonal FLCs) and C57/B6 mice were also injected with the same amount of fibrils or soluble proteins and euthanized at different timepoints for histological analysis.

### Amyloid fibrils purification from organs

For the detection of fibril-associated proteins, amyloid fibrils were purified from mice’s frozen hearts accordingly to previous stablished protocols, with minor modifications (*50*). Briefly, frozen hearts were cut into 1mm pieces, grinded on a 20µm strainer and homogenized on TC buffer (20mM Tris, 138 mM NaCl, 2mM CaCl_2_, pH 8.0) at 4°C with a Kontes pellet pestle homogenizer (DWK Life Sciences). After centrifugation at 3100g, 1 minute at 4 °C, the pellet was washed with ice-cold TC buffer and centrifuged several times, before the incubation in a cocktail of collagenase from *C. histolyticum* (Sigma-Aldrich) and DNAse I (Roche) for 2h at 37°C and 750 rpm. After centrifugation (3100g for 30 min), the pellet was washed 5 times with ice-cold TE buffer (20mM Tris, 140 mM NaCl, 10 mM EDTA, pH 8.0). The resulting pellet was resolubilized with the Kontes pellet pestle homogenizer in ice-cold water and centrifuged at 3100g 5’ at 4°C at least 8 times. Supernatant fractions, as well as the remaining pellet, were analyzed by SDS-PAGE.

For the LC fragmentation analysis, protein deposits from unfixed mice heart tissue were enriched as previously described, with minor modifications (*15*). All procedures were performed on ice using ice-cold buffers. Briefly, tissue specimens were minced with a scalpel, washed 3 times with 1 ml of PBS containing protein inhibitors (Complete, Roche, Basel) and then manually homogenized in 250 µl of Tris EDTA buffer (20 mM Tris, 140 mM NaCl, Complete protein inhibitors, pH 8.0) with a potter pestle. The homogenate was centrifuged for 5 minutes at 3,100 g at 4 °C and the pellet was retained. This step was repeated 10 times overall to remove proteins soluble in the saline solution. The final pellet, enriched in amyloid fibrils and other insoluble proteins, was retained for solubilization and analysis.

### Detection of fibril-associated proteins by MS

20µg of fibril proteins were incubated in 8M urea (10-fold dilution) for 2 hours, then reduced, alkylated and digested with trypsin by filter-assisted sample preparation method. 250 ng of resulting peptides were analysed on a nanoElute2 nano-HPLC system (Bruker Daltonics) that was coupled to a TimsTOF Pro 2 mass spectrometer equipped with CaptiveSpray source (Bruker Daltonics). Peptides were separated on a PepSep C18 column (25cmx150 μm,1.5μm (Extreme) Bruker Daltonics) by a linear 30 min water-acetonitrile gradient from 2% (v/v) to 25% (v/v) of ACN. The timsTOF Pro 2 was operated in DDA-PASEF mode (1.1 sec standard method) using Compass Hystar 6.2. raw data were analyzed by Proteoscape software using the human swissprot database containing the specific immunoglobulin sequence.

### Two-dimensional polyacrylamide gel electrophoresis (2D-PAGE) and western blotting

Pellets were incubated for 2 h with 8 M urea, 4% CHAPS, 0.1 M DTT, followed by protein quantification in supernatants using BCA assay (Thermo Fisher Scientific, Waltham, MA, USA). 15 μg proteins were analyzed by SDS-PAGE on 4-20% acrylamide gradient gels (Biorad), under reducing conditions. For 2D-PAGE analysis, proteins (120 μg for Commasie staining; 50 μg for western blotting) were diluted in Destreak Solution (Cytiva, Merck) plus 0.02% v/v pI 3–10 ampholytes (Bio-Rad). IEF was performed using 11 cm strips, non–linear 3-10 pH gradient (Bio-Rad), followed by SDS-PAGE separation on 8-16% polyacrylamide gradient midi gels (Criterion TGX gels, Bio-Rad), as described. IPG strips were rehydrated at 50 V for 12 h, followed by IEF in a Bio-Rad ProteanTM IEF cell. Proteins were reduced (DTT 0.1 M) and alkylated (IAA 0.15 M) between first and second dimension. Gels were stained with colloidal Coomassie blue (Pierce, Thermo Fisher Scientific, Waltham, MA, USA). For Western blotting, proteins were transferred onto a PVDF membrane (Bio-Rad) using a trans Blot Turbo apparatus (Bio-Rad) and probed with polyclonal rabbit anti-human λ LCs (Dako, Agilent, Santa Clara, CA, USA) used at a concentration of 1 μg/ml, followed by incubation with a horseradish-peroxidase conjugated swine anti-rabbit secondary antibody (Dako). Imaging was performed with an ImageQuant LAS 4000 apparatus (GE Healthcare).

### Analysis of protein spots using liquid chromatography tandem mass spectrometry (LC-MS/MS)

Protein spot excision and in-gel digestion were performed as described (*66*). Peptides extracted from each spot were analyzed by tandem mass spectrometry using an Ultimate 3000 nanoLC system combined with Thermo Scientific™ Q-Exactive Plus Orbitrap mass spectrometer. Tryptic digests were first loaded on a Thermo Scientific Acclaim PepMap C18 cartridge (0.3 mm x 5 mm, 5 μm/100 Å) and then chromatographed on a Thermo Scientific Easy-Spray Acclaim PepMap C18 column (75μm x 15cm, 3μm/100Å packing). MS data were processed using the Mascot software (Matrix Science, London, UK) searching into Swiss-Prot Mus Musculus database, augmented with the sequence of Immunoglobulin lambda light chain. Search parameters were: precursor mass tolerance of 10 ppm, 0.6 Da for HCD fragments, semi-trypsin as the proteolytic enzyme, two missed cleavages, included charge states +2, +3 and +4 and a significance level at p<0.05. Carbamidomethyl (C) was included as fixed modification, Gln->pyro-Glu (N-term Q), Oxidation (M), Pyro-carbamidomethyl (N-term C) were included as variable modifications.

### High resolution ultrasound to evaluate cardiac function of mice

High-resolution data were acquired using the Vevo 3100 high-frequency ultrasound system (FUJIFILM VisualSonics, Canada), equipped with a 40-MHz center frequency MX550D probe, to evaluate left ventricular systolic and diastolic function. Mice were anesthetized with 2.5% isoflurane at 1 ml/min air, and their body temperature was maintained at 37°C using a heating pad. The mice were secured on a stage for imaging, with the ventral thorax hair removed. Throughout the imaging process, the electrocardiogram and respiratory rate were monitored.

We performed 2D echocardiography according to the American Physiological Society guidelines for cardiac measurements in mice (*79*). Longitudinal strain was analyzed in the parasternal long-axis (PLAX) view to obtain left ventricular (LV) ejection fraction and LV longitudinal strain and strain rate at early diastole (SRe). A four-chamber view in B-mode was obtained to evaluate diastolic dysfunction using pulse wave (PW) Doppler to assess isovolumic relaxation time, E/A ratio, and E/E’ ratio. A short axis view was used to assess anterior and posterior wall thickness in diastole and systole. LV mass corrected was calculated using this formula: 0.8424 x [(LVID;d + LVPW;d + IVS;d)^3^ – LVID;d^3^]. To assess left atrium anatomy and function, we used 4D ultrasound (US) imaging. The US probe was attached to a linear step motor scanning in the parasternal short-axis (PSAX) view from below the apex to above the aortic arch. Acquisition parameters were set to a gain of 48 dB, 3D range of 12-15 mm, 3D step size of 0.127 mm, and frame rate of 400 frames/sec. Each image acquisition required 10–15 minutes to create 4D data throughout one representative cardiac cycle from base to apex. We evaluated the atrial volume at systole and diastole and calculated the left atrium (LA) ejection fraction using the formula: stroke volume / end diastolic volume.

These three measurements provide insights into the heart’s filling and relaxation. Offline image analysis was conducted using Vevo Lab Software 5.8.1 (FUJIFILM VisualSonics).

### Transcriptional analysis and RNA-seq

Total RNA was extracted from the apical part of the hearts of λS-DH and DH-LMP2A mice using the miRNeasy Mini Kit (Qiagen) on the TRIzol reagent and the purity of the samples was assessed on the Bioanalyzer RNA 6000 Nano (Agilent) before generating the libraries and performing the RNA-sequencing as previously described (*77*). Paired-end reads were processed through the bioinformatic pipeline nf-core/rnaseq v3.12 (*80*) in reverse strandedness mode. This performed, quality control, alignment to the GRCm38 mouse genome with STAR (*81*) and quantification by Salmon(*82*) with the corresponding genome annotation from the ENSEMBL release 111 database. The batch effect due to the two separate sequencing run was taken into account through ComBat adjustment (*83*) and the differential analysis performed by EdgeR (Bioconductor) (*84*). Genes were considered differentially expressed (DE) if they had a corrected p-value (FDR) ≤ 0.05 and an absolute foldchange ≥ 1.5. GSEA analysis of the Reactome, Gene Ontology (GO) Biological Process, Cellular Components and Molecular Function enriched pathways, as well as cell type enrichment by the Mouse Cell Atlas, were performed on WEB-based GEne SeT AnaLysis Toolkit (Web-Gestalt) (*85*) using the list of DE genes.

### Analysis of the clearance of LCs

C57/B6 mice were injected intravenously with 1 mg of λS-LCs. Plasma was taken after the injection, and at several timepoints (30 minutes, 1h, 3h and 6h) to asses the clearance of the LCs in the mice. Dosage of the LCs was performed by ELISA with the appropriated antibody.

### In silico study of the **λ**S-VL sequence

The aggregation prediction was generated on Aggrescan4D (https://biocomp.chem.uw.edu.pl/a4d/) (*41*) by using the already published structure from the recombinant 6aJL2 germline IGLV6-57 sequence (PDB: 2W0K) (*86, 87*). The germline sequence was mutated to obtain the sequence from the λS-VL, and the resulting structures were analyzed at physiological pH (7.5) at an interaction distance of 5 Å.

## ACKNOWLEDGMENTS

The authors thank the staff of the Biologie Intégrative Santé Chimie Environnement (BISCEm) technical platforms at the University of Limoges (animal, cell cytometry and microscopy transgenesis facilities), the Department of Pathology of Poitiers. The authors also thank the GeT-Santé facility (I2MC, Inserm, Génome et Transcriptome, GenoToul, Toulouse, France) for the advice and technical contribution to the experiments of the RNA sequencing experiments.

This work was supported by grants from Agence National de la Recherche (#ANR-21-CE17-0040-01), Fondation pour la Recherche Médicale (# FRM-EQU202203014615), Fondation Franc_Jaise pour la recherche sur le Myelome et les Gammapathies monoclonales (FFRMG) and Ligue nationale contre le cancer. GM-R is funded by fellowships from Région Nouvelle Aquitaine and Société Française d’Hématologie. MVA was funded by fellowship from Fondation ARC pour la Recherche sur le Cancer. SB is supported by Centre Hospitalier Universitaire Dupuytren Limoges and Plan National Maladies Rares.

