## Supplemental Figures for "A mouse model of cardiac AL amyloidosis unveils mechanisms of tissue accumulation and toxicity of amyloid fibrils"

**Supplemental figure 1: Characterization of the λS-LC**

A. Amino acid sequence alignment between the λS-LC (LS) and the germline sequence IGLV6-57 according to the IMGT sequence. Other LCs from the same family were also aligned to our sequence (Wil, Jto and AL55). Mutated amino acids are indicated in yellow, with the percentage of shared sequence is indicated for each one of the alignments.
B. *In silico* analysis of the structure of λS-VL sequence with Aggrescan4D software (https://biocomp.chem.uw.edu.pl/a4d/) showed three mutations towards hydrophobic amino acids (Ile29, Ile65 and Phe80) that were absent in the germinal sequence Vλ6-57 (PDB: 2W0K). The green squares correspond to the Asp1 (N-terminal). The aggregation score is given at pH 7.5.
C. Kinetics of the clearance of λS-LC injected in C57/B6 mice (n=3) followed by ELISA in plasma 30 minutes, 1 hour, 3 hours and 6 hours after intravenous injection of 1 mg of protein showed the elimination of % of the protein within the first hour.
D. Confirmation of the presence of the λS-LCs in the serum of our λS-DH transgenic mice by Western Blot with an antibody recognizing the human Vλ6-57 LCs, and the absence of this human LC in the serum of controls DH-LMP2A (DH) and κR-DH mice.
E. Physiological detection of human λ -LCs in the spleen and the kidney of λS-DH mice, consistent with the production and the reabsorption of LCs, respectively.

**Supplemental figure 2: Analysis of recombinant λS proteins**

A. SDS-PAGE of the purified recombinant λS proteins, the full-length LC (rλS-LC, ~25kDa) and its variable domain (rλS-VL, ~12kDa) under reducing and non-reducing conditions. Total protein was visualized by the StainFree technology (Biorad).
B. Size exclusion chromatography of the rλS-VL confirmed that there were no aggregated forms of the rλS-VL in our samples. Lysozyme (~16.5kDa) and albumin (~66kDa) were used as size standards.

**Supplemental figure 3: Cardiac amyloid deposits in λS-DH mice**

A. Representative images of the amyloidosis score in the heart of the λS-DH mice according to the CR-stained deposits in fluorescence. “No”, or score 0, corresponds to the total absence of CR staining. “Low”, or score 1, corresponds to one of some focal deposits within the myocardium or/and blood vessels. “Mid”, or score 2, corresponds to some diffuse deposits within the myocardium. “High”, or score 3, corresponds to the diffuse deposits throughout the myocardium.
B. CR and the hλ-LC fluorescence in the heart of low amyloid score λS-DH mice, are colocalized, but more extended hλ-LC can be observed in the intercellular spaces surrounding the CR positive areas.
C. CR and the hλ-LC staining in mice tissues did not show any amyloid-related staining in the control mice induced with the same protocols as λS-DH mice. Example of a DH mouse 7 days after the injection of seeds.

**Supplemental figure 4: Amyloid deposition in other tissues in λS-DH mice**

CR staining revealed that AL amyloid deposits could also be found in other tissues in our λS-DH mice, such as the tongue, liver, fat and lung. Example of a mid-score mouse 4 months after induction with the seeding protocol.

**Supplemental figure 5: Rare spontaneous amyloidosis in the λS-DH mice**

A. Spontaneous amyloidosis occurring in the 28-mice cohort analyzed after 12 months. Two mice from the cohort developed AL amyloidosis spontaneously at 14 and 18 months. The cardiac score of these mice was evaluated as mid.

**Supplemental figure 6: LC fragmentation patterns in λS-DH micen control mice and human**

A. Western blotting analysis with a primary antibody against the hλ–LCs of the insoluble material enriched from the heart of amyloid-positive and amyloid-negative mice. For the group of amyloid-positive mice, only a representative positive sample is shown.
B. 2D western blotting analysis on *ex vivo* fibril-constituting LCs from fat (F) and cardiac (H) tissue extracted from an AL patient with an IGLV6-57 LC (AL55). The bottom panel shows the boxed region of the top panel. Adapted with permission from Mazzini et al, FEBS, 2021 *(1)*

**Supplemental figure 7: Echocardiography analysis**

Cardiac scores from the AL+ λS-DH mice analyzed by plasmatic NT-proBNP dosage (mean 2.8, range =1, n=10).

A. 4DUS image of the left atrium appendage thrombus in the λS-DH mouse excluded from the analysis
B. Cardiac scores from the AL+ λS-DH mice analyzed by high resolution echocardiography (mean 2.5, range =2, n=6).
C. Calculated left ventricular (LV) mass from control DH (Sham) and AL+ λS-DH mice (AL). A non-significant 37% mean increase was observed in AL group compared to controls (p=0.0649)
D. Posterior LV thickness measured in systole (s) showed a non-significant 37% increase in the AL mice (p=0.0606). Anterior LV thickness measured in systole and diastole (d) showed an increase of 23% (p=0.0108) and non-significant 27% (p=0.0649) respectively in AL mice compared to controls.
E. Calculated LV ejection fraction calculated by the longitudinal strain analyzed in parasternal long-axis view did not show a significant difference between the control and the AL+ λS-DH mice (p=0.8182).
F. Global LV longitudinal peak strain analyzed in parasternal long-axis view and early to late diastolic transmitral flow velocity was given by the ratio E/A did not show any significant difference (p=0.8182 and 0.6991 respectively).
G. Left atrium (LA) volume evaluated at systole and the calculated left atrial ejection fraction showed a 1.7 fold increase (p=0.0152) and 1.7 fold decrease (p=0.0087) respectively.

**Supplemental figure 8: Transcriptomic analysis of hearts from λS-DH mice and controls**

A. Cardiac scores from the AL+ λS-DH mice analyzed by RNA sequencing (n=10).
B. Principal component analysis showing the distribution of the AL+ λS-DH (AL), AL- λS-DH (LS) and DH (Ctrl) samples analyzed by RNA sequencing according to the variance in gene expression.
C. Heatmaps showing the 3887 DEG among the 10059 genes analyzed in the comparison between λS-DH mice with amyloid deposits (AL) and both λS-DH and DH mice without amyloid deposits (LS and Ctrl). Mice were clustered according to the DE profiles. Red corresponds to overexpressed genes and blue to downregulated genes.
D. Reactome enriched pathways of the 3887 DEG from this analysis. The represented pathways were significantly enriched (FDR < 0.05)
E. Gene Ontology analysis of the DEG according to the biological process (top left), cellular components (bottom left) and molecular function (right). The represented processes were significantly enriched (FDR < 0.05)
F. Mouse Cell Atlas analysis showing the cell type enrichment in the cardiac tissue according to the DEG. The represented cell types were significantly enriched (FDR < 0.05)
A. Gene Set Enrichment plot representing the distribution of the 55 DEG related to the extracellular matrix organization reactome pathway.

### Fig. S1

## A.

```
              FR1-IMGT                  CDR1-IMGT        FR2-IMGT        CDR2-IMGT                 FR3-IMGT                     CDR3-IMGT     FR4-IMGT
               (1-26)                    (27-38)          (39-55)         (56-65)                  (66-104)                     (105-117)     (118-128)

         1        10        20         30          40        50          60          70        80        90        100
         .........|.........|......   ...|........ .|.........|..... ....|.....  ....|.........|.........|.........|....  .....|.........|.........
LS       DFMLTQPHS.VSESPGKTVTISCTRS SGIL....ASNS VQWFQQRPGSSPTTVIY GG.......I QRPSGVP.DRFSGSFDSSSNSASLSISGLKTEDEADYYC QSYDS...TNEGV FGGGTKLTVL
IGLV6-57 NFMLTQPHS.VSESPGKTVTISCTRS SGSI....ASNY VQWYQQRPGSSPTTVIY ED.......N QRPSGVP.DRFSGSIDSSSNSASLTISGLKTEDEADYYC QSYDS....SNVV FGGGTKLTVL (86.4%)
         D                            IL      S    F              GG       I               F         S                           TNEG

Wil      NFLLTQPHS.VSESPGKTVTISCTRS SGSI....ANNY VHWYQQRPGSSPTTVIF ED.......D HRPSGVP.DRFSGSVDTSSNSASLTISGLKTEDEADYYC QSYDH....NNQV FGGGTKLTVL (80.0%)
Jto      NFMLNQPHS.VSESPGKTVTISCTRS SGNI....DSNY VQWYQQRPGSAPITVIY ED...... N QRPSGVP.DRFAGSIDRSSNSASLTISGLKTEDEADYYC.QSYDA....RNVV FGGGTRLTVL (79,1%)
AL55     NFMLTQPHS.VSESPGKTLTISCTGS SASI....ASHY VQWYQQRPGGAPTTLIY EN.......D QRPSEVP.DRFSGSIDSSSNSASLTISGLKTEDEADYYC.QSYDG...NNHWV FGGGTKLTVL (80,2%)
```

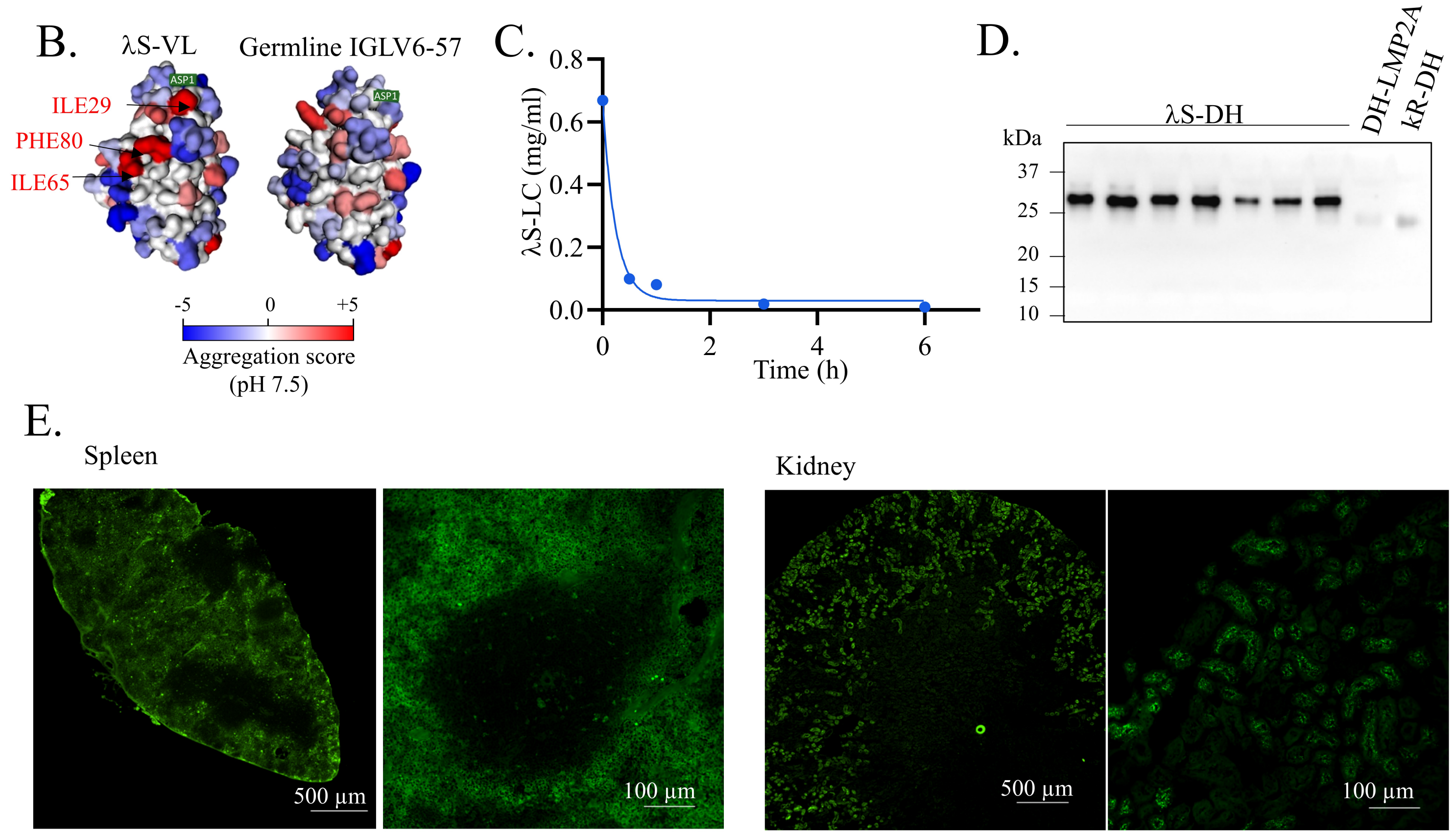

### Fig. S2

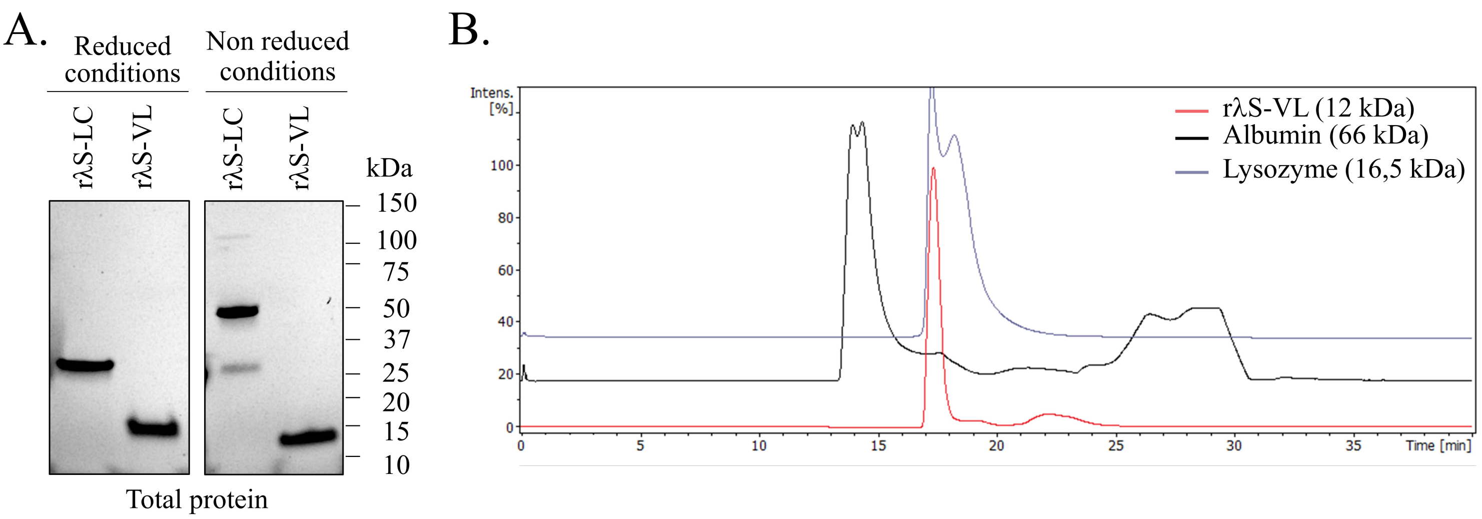
A.
Reduced conditions
Non reduced conditions
rλS-LC
rλS-VL
rλS-LC
rλS-VL
kDa
150
100
75
50
37
25
20
15
10
Total protein
B.
Intens. [%]
100
80
60
40
20
0
0
5
10
15
20
25
30
35
Time [min]
rλS-VL (12 kDa)
Albumin (66 kDa)
Lysozyme (16,5 kDa)

#### Fig. S3

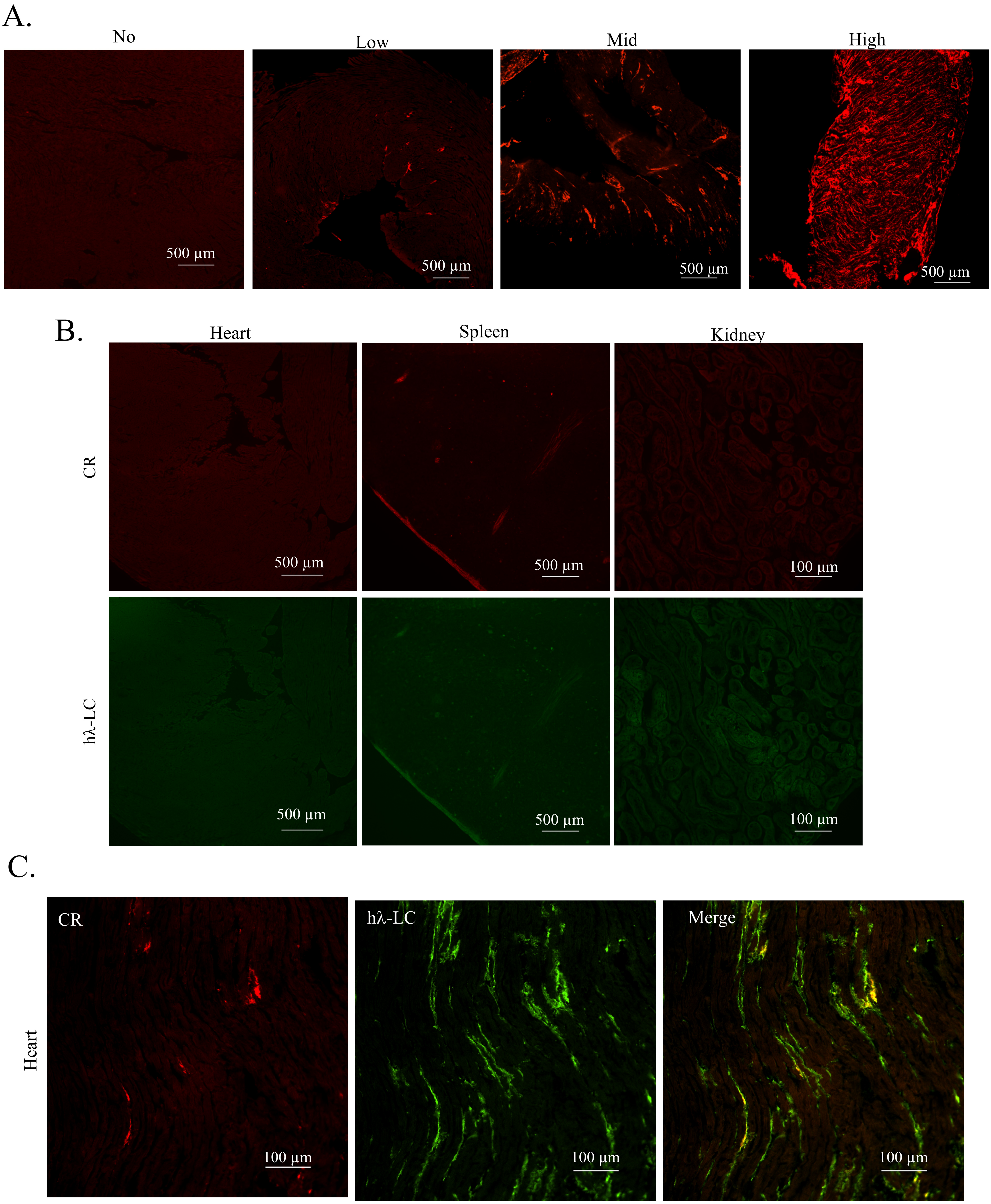
A.
No
Low
Mid
High
500 µm
500 µm
500 µm
500 µm
B.
Heart
Spleen
Kidney
CR
hλ-LC
500 µm
500 µm
100 µm
500 µm
500 µm
100 µm
C.
CR
hλ-LC
Merge
Heart
100 µm
100 µm
100 µm

**Fig. S4**

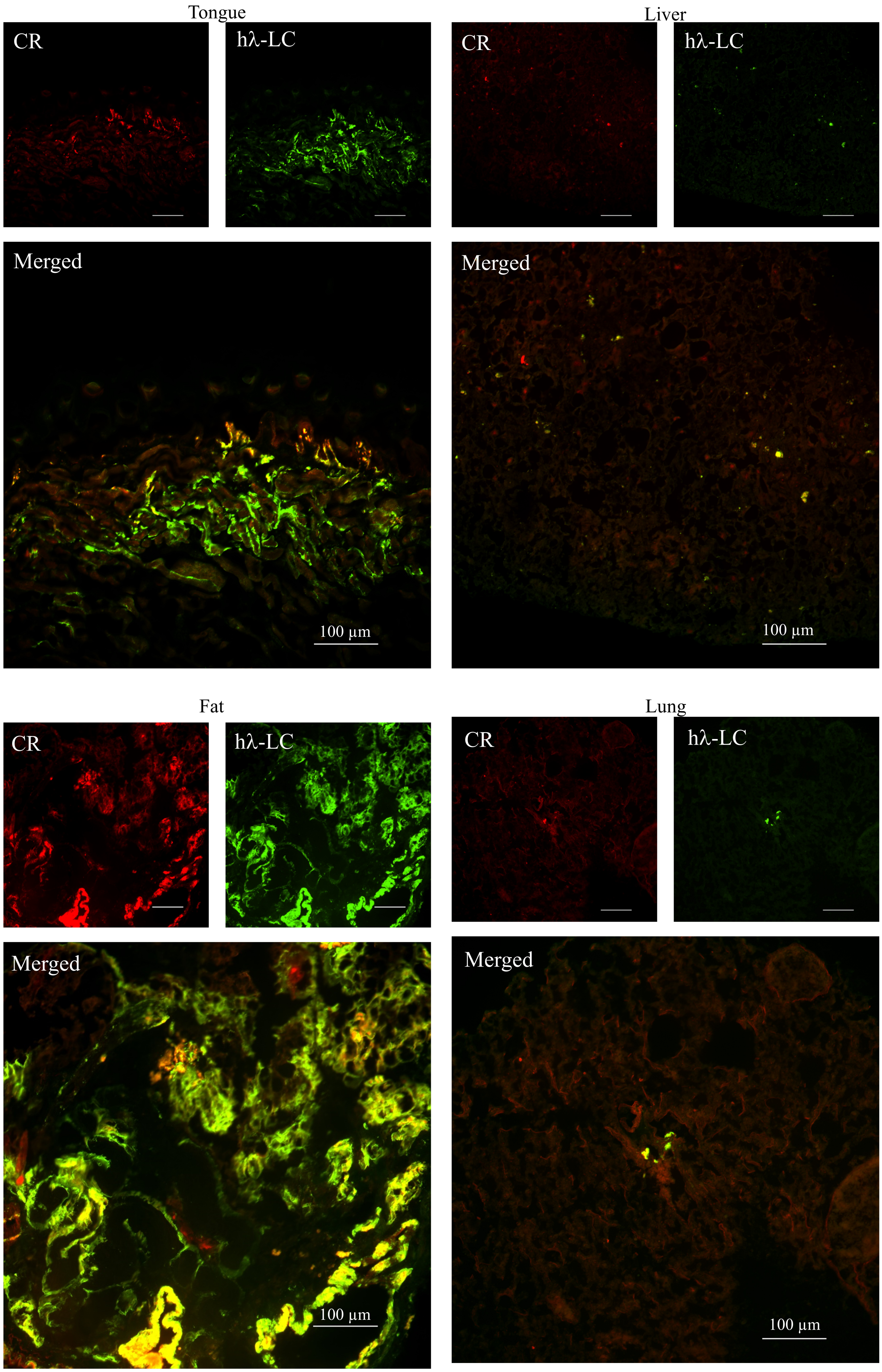
Tongue
CR
hλ-LC
Merged
100 μm
Liver
CR
hλ-LC
Merged
100 μm
Fat
CR
hλ-LC
Merged
100 μm
Lung
CR
hλ-LC
Merged
100 μm

### Fig. S5

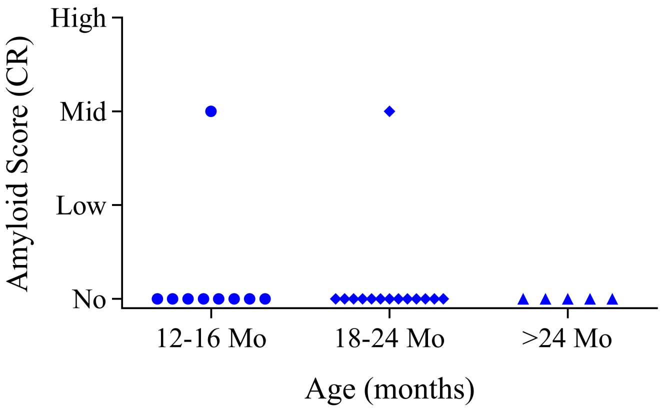

High
Mid
Low
No
Amyloid Score (CR)
12-16 Mo
18-24 Mo
>24 Mo
Age (months)

### Fig. S6

A.

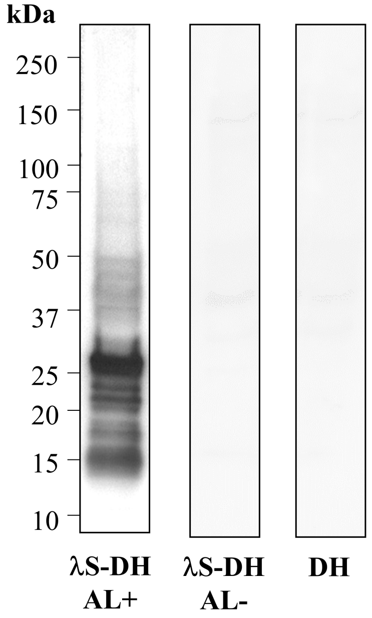
kDa
250
150
100
75
50
37
25
20
15
10
λS-DH AL+
λS-DH AL-
DH

B.

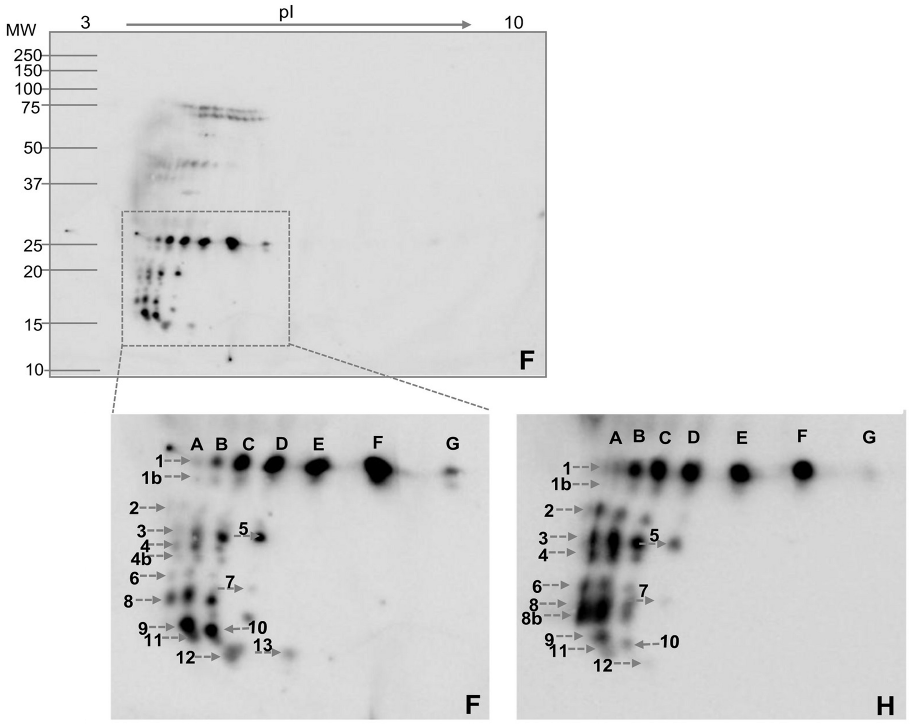
MW
3
pI
10
250
150
100
75
50
37
25
20
15
10
F
A B C D E F G
1
1b
2
3
4
4b
5
6
7
8
9
10
11
12
13
F
A B C D E F G
1
1b
2
3
4
5
6
7
8
8b
9
10
11
12
H

### Fig. S7

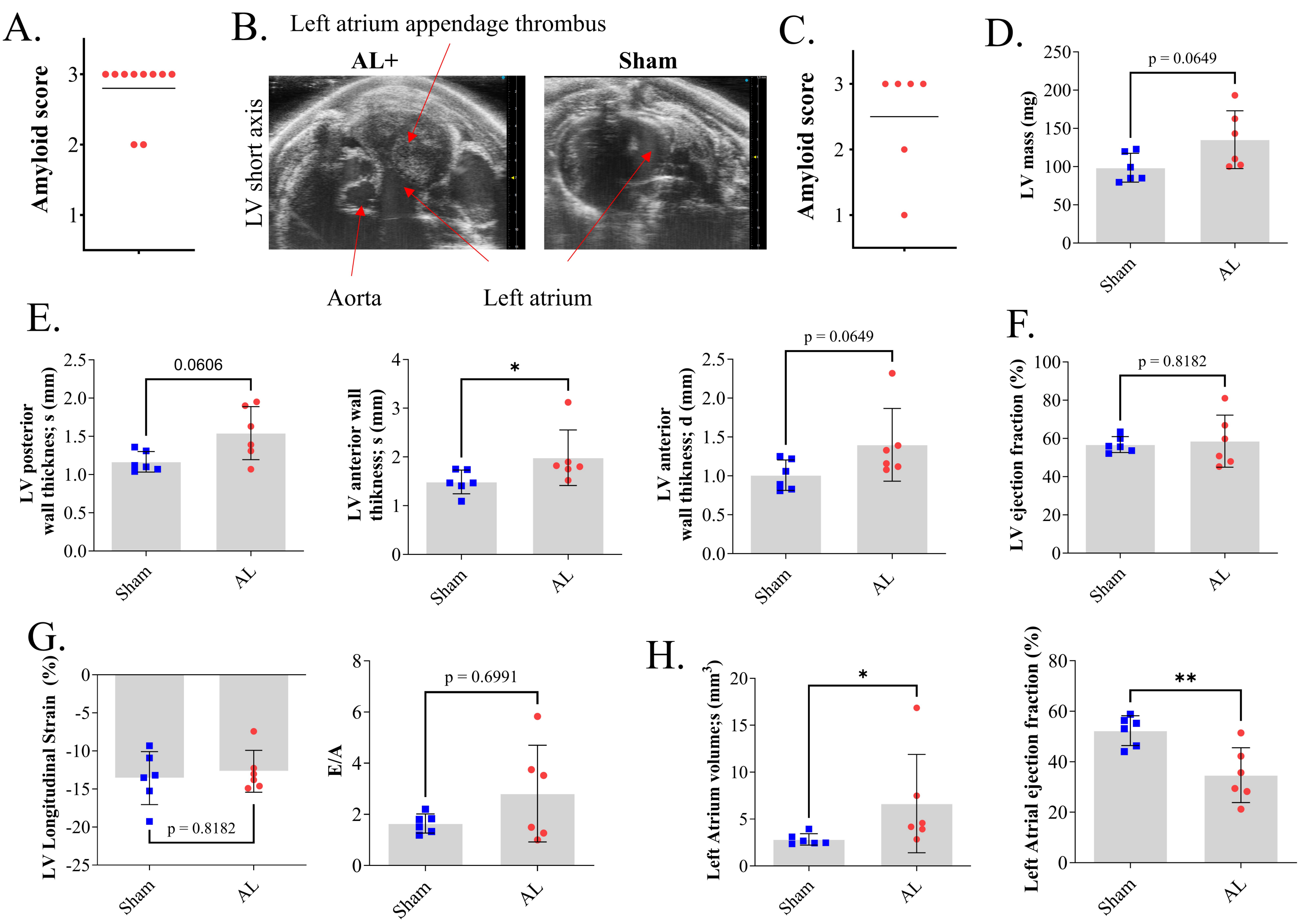
A.
Amyloid score
B.
Left atrium appendage thrombus
AL+
Sham
LV short axis
Aorta
Left atrium
C.
Amyloid score
D.
LV mass (mg)
p = 0.0649
Sham
AL
E.
LV posterior wall thickness; s (mm)
0.0606
LV anterior wall thikness; s (mm)
*
LV anterior wall thikness; d (mm)
p = 0.0649
F.
LV ejection fraction (%)
p = 0.8182
G.
LV Longitudinal Strain (%)
p = 0.8182
E/A
p = 0.6991
H.
Left Atrium volumes (mm3)
*
Left Atrial ejection fraction (%)
**

### Fig. S8

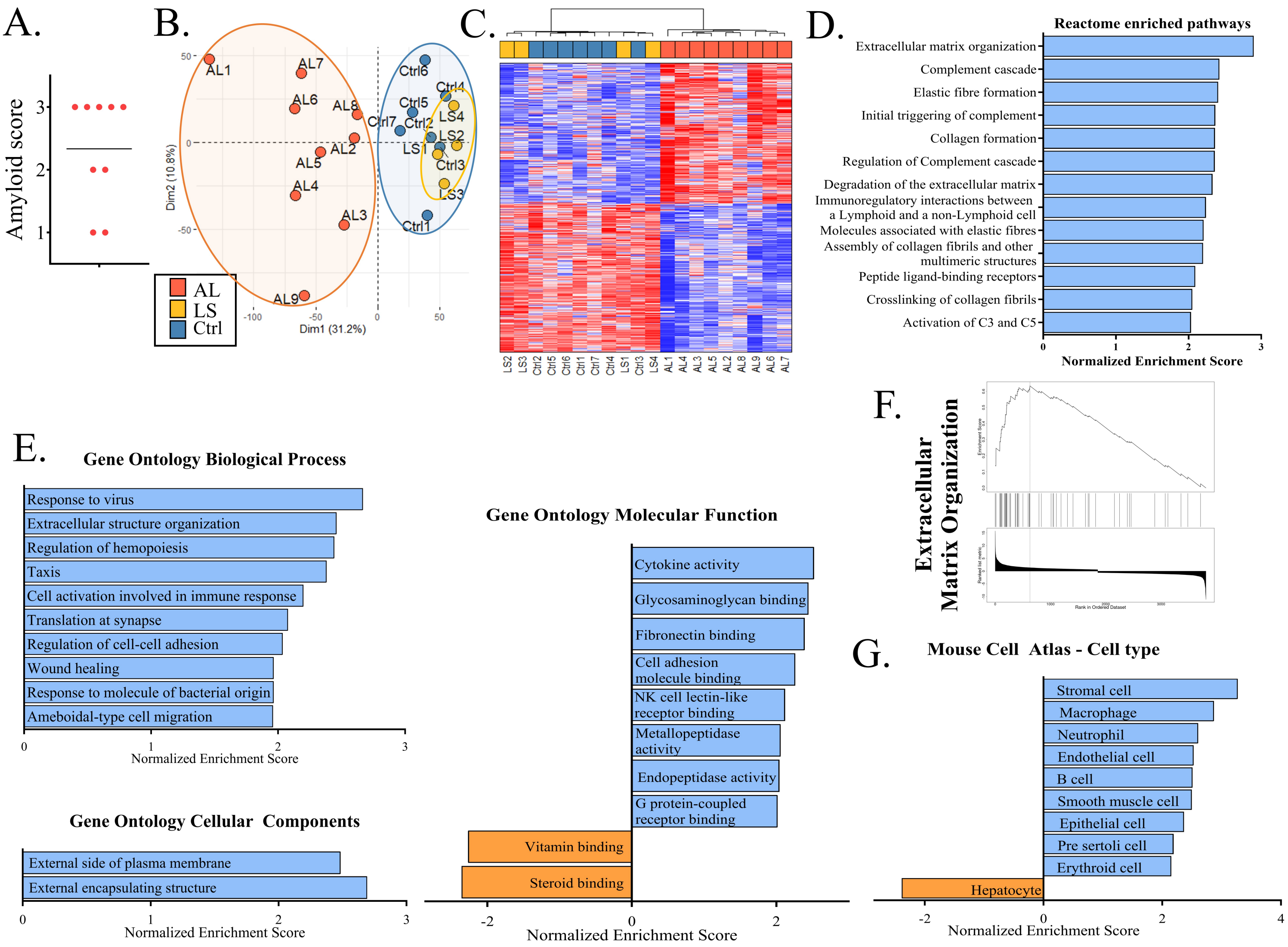
A.
Amyloid score
3
2
1
B.
AL1
AL7
AL6
AL8
AL2
AL5
AL4
AL3
AL9
Ctrl6
Ctrl4
Ctrl5
Ctrl7
Ctrl2
LS4
LS2
LS1
Ctrl3
LS3
Ctrl1
Dim2 (10.8%)
Dim1 (31.2%)
50
0
-50
-100
AL
LS
Ctrl
C.
LS2
LS3
Ctrl2
Ctrl5
Ctrl6
Ctrl1
Ctrl7
Ctrl4
LS1
Ctrl3
LS4
AL1
AL4
AL3
AL5
AL2
AL8
AL9
AL6
AL7
D.
Reactome enriched pathways
Extracellular matrix organization
Complement cascade
Elastic fibre formation
Initial triggering of complement
Collagen formation
Regulation of Complement cascade
Degradation of the extracellular matrix
Immunoregulatory interactions between a Lymphoid and a non-Lymphoid cell
Molecules associated with elastic fibres
Assembly of collagen fibrils and other multimeric structures
Peptide ligand-binding receptors
Crosslinking of collagen fibrils
Activation of C3 and C5
0
1
2
3
Normalized Enrichment Score
E.
Gene Ontology Biological Process
Response to virus
Extracellular structure organization
Regulation of hemopoiesis
Taxis
Cell activation involved in immune response
Translation at synapse
Regulation of cell-cell adhesion
Wound healing
Response to molecule of bacterial origin
Ameboidal-type cell migration
0
1
2
3
Normalized Enrichment Score
Gene Ontology Cellular Components
External side of plasma membrane
External encapsulating structure
0
1
2
3
Normalized Enrichment Score
Gene Ontology Molecular Function
Cytokine activity
Glycosaminoglycan binding
Fibronectin binding
Cell adhesion molecule binding
NK cell lectin-like receptor binding
Metallopeptidase activity
Endopeptidase activity
G protein-coupled receptor binding
Vitamin binding
Steroid binding
-2
0
2
Normalized Enrichment Score
F.
Extracellular Matrix Organization
Rank in Ordered Dataset
G.
Mouse Cell Atlas - Cell type
Stromal cell
Macrophage
Neutrophil
Endothelial cell
B cell
Smooth muscle cell
Epithelial cell
Pre sertoli cell
Erythroid cell
Hepatocyte
-2
0
2
4
Normalized Enrichment Score

### Table S1

| Gene | Log FC | FDR | Description |
|---|---|---|---|
| Osbpl3 | 1.72 | 0.0007 | Encodes one of the members of the Oxysterol-Binding protein family, an intracellular lipid receptor family involved in the maintenance of cholesterol levels in the organism |
| Snx10 | -1.17 | 0.0083 | Encodes a member of the Sortin Nexin family, involved in intracellular trafficking in endosomes through their binding with phosphoinositide |
| Pyurf | -1.77 | 0.0023 | Encodes for the PIGY Upstream Open Reading Frame, an S-adenosylmethionine-dependent methyltransferase chaperone involved in the stabilization of the coenzyme Q components during its biosynthesis in mitochondria. |

### Table S2

| | **ALvsCtrl** | | **ALvsLS** | |
|---|---|---|---|---|
| | **Log2 FC** | **FDR** | **Log2 FC** | **FDR** |
| Nmrk2 | 3,91 | 0,0007 | 3,44 | 0.0062 |
| Irf7 | 2,09 | 0,0033 | 3.45 | 0,0011 |
| Gpnmb | 4,12 | 1,85E-05 | 3.47 | 0,0001 |
| Mmp12 | 7,6 | 0,0026 | 8,17 | 0,0098 |
| Lgals3 | 2,7 | 0,0001 | 4,09 | 0,0002 |
| Ltbp2 | 2,71 | 0,0029 | 3,52 | 0,0058 |
| Timp1 | 2,63 | 0,0094 | 3,74 | 0,0129 |
| Nppa | 3,58 | 0,0021 | 1,992 | 0,095 |

### Table S3

| Antibody | Reactivity | Clone | Source | Conjugate | Applications |
|---|---|---|---|---|---|
| Anti-B220 | Mouse | RA3-6B2 (Becton Dickinson) | Rat | BV421 | Flow cytometry |
| Anti-CD138 | Mouse | 281-2 (BD Biosciences) | Rat | APC | Flow cytometry |
| Anti-human VL6 | Human | Monoclonal (from A. Solomon’s lab) | Mouse | Unconjugated | Western-Blot |
| Anti-lambda | Human | Polyclonal (southernBiotech) | Goat | Unconjugated or AP | ELISA |
| Anti-lambda | Human | Polyclonal (Beckman Coulter) | Goat | Unconjugated, AP, HRP, gold particules | ELISA |
| Anti-lambda | Human | Polyclonal (Dako) | Rabbit (Fab'2) | FITC | Flow cytometry+ IF + Western-Blot |
| Anti-lambda | Human | [4C2] Monoclonal | Mouse | FITC | IF |
| Anti-IgG | Rabbit | Polyclonal (Beckman Coulter) | Goat | HRP | Western-Blot |
| Anti-IgG | Mouse | Polyclonal (Beckman Coulter) | Goat | HRP | Western-Blot |
